# Metabolic buffering suppresses phenotype switching in cancer

**DOI:** 10.1101/2025.04.07.647576

**Authors:** Ana Ramírez-Sánchez, Miguel Jociles-Ortega, José Manuel García-Martinez, Irene Torrens-Martínez, Javier Martínez-Useros, María Jesús Fernández-Aceñero, Teresa Olmos-De Blas, Pakavarin Louphrasitthiphol, Colin R. Goding, Custodia García-Jiménez, Ana Chocarro-Calvo

## Abstract

The impact of the microenvironment on epigenetically plastic cancer cells underpins phenotypic heterogeneity, a major cause of metastatic dissemination and therapy resistance that together represent the primary cause of cancer-related death. Nutrient limitation is a key microenvironmental stress that can cause a phenotypic transition from proliferation to invasion via activation of the integrated stress response. However, whether and how the capacity to store and mobilize nutrients impacts phenotype-switching through metabolic buffering remains unknown. Here, using melanoma as a model, we reveal that the ability to accumulate and mobilize glycogen, that buffers glucose availability, plays a key role in phenotypic transitions in melanoma. While proliferative phenotype cells exhibit high levels of glycogen, invasion is marked by low glycogen levels. Significantly, an inability to store and metabolize glycogen leads to phenotype instability and a switch to invasion. Accordingly, glycogen levels inversely correlate with Clark levels in primary melanomas, with low expression of the glycogen phosphorylases PYGB/L and phosphoglucomutase 1 (PGM1) being associated with worse overall survival. The importance of metabolic buffering in suppressing phenotypic transitions likely extrapolates to other cancer types.

**Highlights:**

- Melanoma phenotypes are distinguished by their ability to store and mobilize glycogen.
- Proliferative MITF^High^ melanoma cells store glycogen to improve survival under stressful conditions.
- Inhibition of glycogen degradation impairs proliferation in MITF^High^ melanoma cells.
- Lack of PGM1 drives invasion and metastatic dissemination.

## Introduction

Cancer cell phenotypic heterogeneity underpins metastasis, contributes to therapy resistance, and poses a major therapeutic challenge. In addition to changes in metabolism and gene expression driven by oncogene activation, in response to a changing microenvironment, cancer cells undergo dynamic and reversible phenotypic transitions. These are accompanied by both metabolic rewiring (1) and remodeling of the epigenetic landscape, a hallmark of malignant transformation across all cancer types (2). Different phenotypic states within a tumor can therefore exhibit fundamentally different metabolic profiles (3). Changes in metabolism and any associated phenotypic transition may in principle be suppressed by the presence of ‘metabolic reservoirs’ that might buffer against perturbations in nutrient availability, one of the primary causes of phenotype switching in cancer (4). The concept of metabolic reservoirs in cancer is relatively new (5) and envisages that energy-producing molecules can be stored by cancer cells for later use. One key metabolic reservoir is glycogen that contributes to glucose homeostasis in cancer cells by supplying glucose for glycolysis and carbons for the pentose phosphate pathway and the tricarboxylic acid cycle (6). However, whether the amount of glycogen stored in a cell or glycogen metabolism in general plays a critical role in phenotype- switching and contributing to the generation intra-tumoral heterogeneity is poorly understood.

Melanoma represents an excellent model to study phenotype-switching. Multiple phenotypic states are well- defined by the expression of key markers including the microphthalmia-associated transcription factor (MITF). MITF controls many features of melanoma biology (7) and promotes either differentiation or proliferation, regulates metabolism and DNA damage repair, and suppresses invasion. Cell intrinsic events such as activation of BRAF, or microenvironmental factors, such as glutamine (8) or glucose limitation (9), down-regulate MITF mRNA and protein expression and lead to a reversible switch from proliferation to invasion. Significantly, the switch from an MITF^High^ to MITF^Low^ state is accompanied by altered metabolism; proliferative phenotype melanoma cells depend primarily on glycolysis (10) whereas invasive melanoma cells, that are also slow- cycling and therapy resistant, obtain their energy primarily through oxidative phosphorylation (11).

Here, using melanoma as a model, we explore whether metabolic buffering, via the accumulation of glucose stores in the form of glycogen, impacts the ability of melanoma to undergo phenotypic transitions. The results reveal that accumulation of glycogen is a hallmark of proliferative, but not invasive phenotype cells, and that low of glycogen stores leads to phenotypic instability and a rapid transition to invasion.

## Results

### Proliferative, MITF^High^, melanoma cells accumulate glycogen and are highly glycolytic

Since proliferating cancer cells have increased demands for glucose and intracellular glycogen stores have been observed in various tumors (12), we set out to determine the role and importance of glycogen storage in melanoma by first characterizing the glycogen content in human tumors. Periodic acid–Schiff stain (PAS) with or without the glycogen-degrading enzyme amylase (PASD), as a control for the specificity of the staining, was performed in patient melanoma samples as well as in nevi. 67 samples were included in total, of which 14 were nevi (20.9%), 26 primary melanomas (38.8%) and 27 metastasis (40.3%) (see Table 1 for patient and tumor characteristics). Nevi were characterized by small glycogen accumulations in the upper layers of the epidermis, evidenced by intense magenta staining that disappeared when pre-digested with diastase (PAS/D) (Fig. 1A, left panels). In contrast, primary melanomas were either PAS-positive (Fig. 1A, middle panels) or negative (Fig. S1A); in PAS positive melanomas, glycogen specific staining was localized throughout the tissue (Fig. 1A). Consistently, metastases showed less glycogen accumulation than primary melanoma samples (Fig. 1A right panels, quantified in Fig. 1B), with a trend towards significance (p=0.094).

**Fig. 1.**
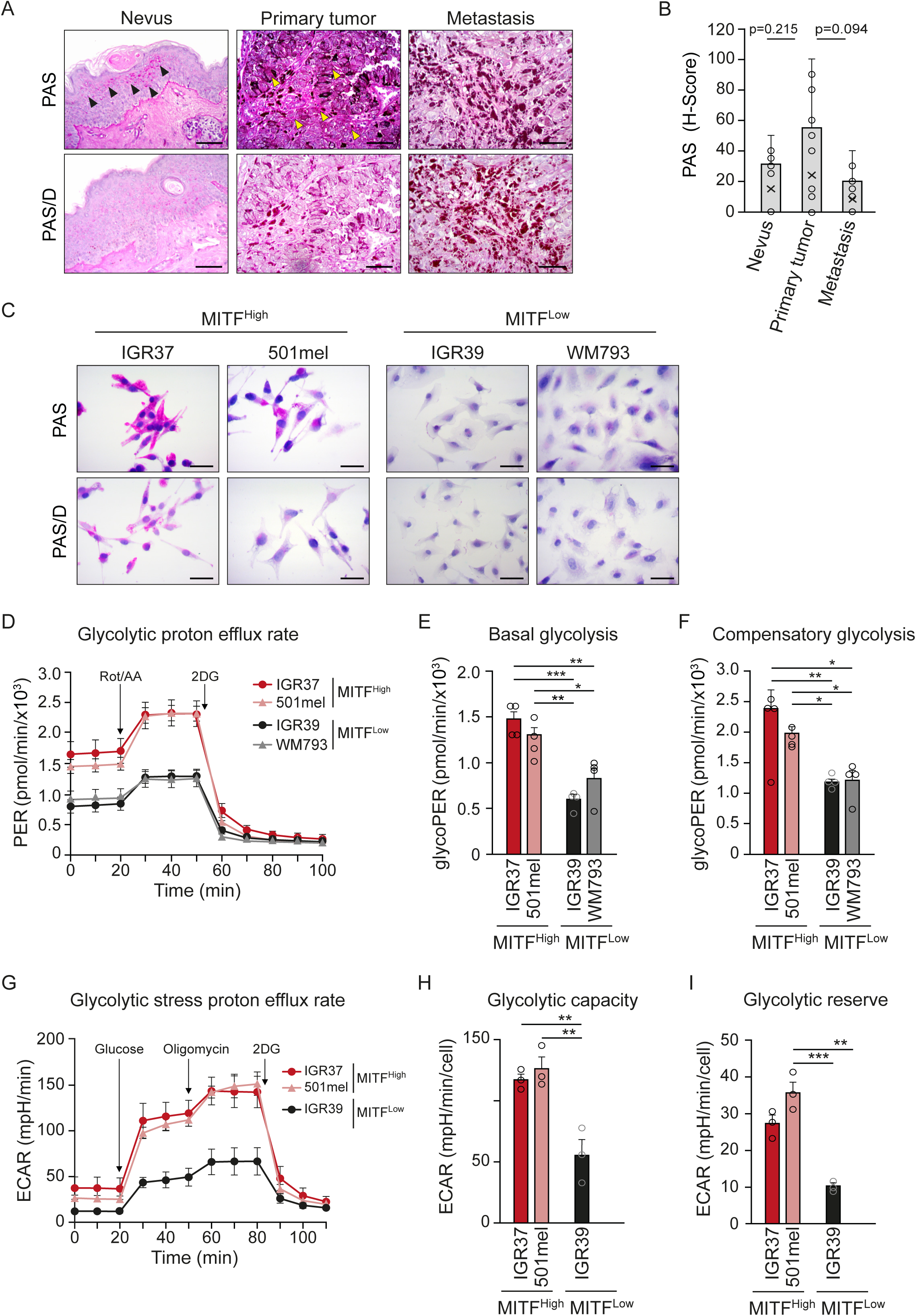
Melanoma MITF^High^ glycolytic cells accumulate glycogen. (A) Immunohistochemical analysis of glycogen content in human melanoma biopsies of nevus (left), melanoma (middle) and metastasis (right) stained with Periodic Acid-Shiff (PAS) (upper) or digested with diastase (PAS/D) (lower) to assess the specific glycogen staining in consecutive sections. Arrows showing glycogen stores localization. Brown colour represents tissue pigmentation (melanin). Scale bars: 50 μm. (B) Quantification of PAS staining from human samples in (A) n =14 for nevus, n=27 for primary melanoma and n=28 for metastasis. The profile was obtained from Histoscore (H-Score) variations. Statistical analysis by Man Whitney U-test; (C) Glycogen stores in IGR37 and 501mel (MITF^High^) and IGR39 and WM793 (MITF^Low^) melanoma cells revealed by PAS staining or treated with diastase (PAS-D). Scale bars: 25 μm. (D) Glycolytic Proton Efflux Rate (PER) of IGR37 and 501mel (MITF^High^) and IGR39 and WM793 (MITF^Low^) melanoma cells. n=4 for IGR37, 501mel and IGR39 cells, n=5 for WM793; error bars indicate SEM (E, F) Quantification of glycolytic parameters derived from (D). Bar graphs representing Basal (E) and Compensatory (F) glycolysis rates. n=4. (G) Extracellular acidification rate (ECAR) of IGR37 and 501mel (MITF^High^) and IGR39 and WM793 (MITF^Low^) melanoma cells. n=3. (H, I) Quantification of glycolytic parameters derived from (G). Bar graphs representing Glycolytic capacity (H) and reserve (I) from 3 experiment. Mean ± SEM of replicates is displayed for panels E-F, H and I. Statistical analysis by One-Way ANOVA was performed; *P < 0.05; **P < 0.01; ***P < 0.001.

**Table 1.** Clinicopathological characteristics of patients.

| Patients Characteristic | N (%) |  |  |
| --- | --- | --- | --- |
|  | Nevus | Primary melanoma | Metastasis |
| <u>Gender</u> |  |  |  |
| Male | 4 (28,6%) | 11 (40,7%) | 15 (53,6%) |
| Female | 10 (71,4%) | 15 (55,6%) | 12 (42,8%) |
| N/A | 0 | 1 (3,7%) | 1 (3,6%) |
| <u>Median Age (years)</u> |  |  |  |
| <u>Melanoma Classification</u> |  |  |  |
| MES |  | 4 (28.6%) |  |
| MN |  | 6 (42.9%) |  |
| LMM |  | 3 (21.4%) |  |
| MLA |  | 1 (7.1%) |  |
| <u>Breslow Index</u> |  |  |  |
| In situ |  | 4 (13.3%) |  |
| Intermediate |  | 11 (36.7%) |  |
| High |  | 15 (50%) |  |
| <u>Clark level</u> |  |  |  |
| I |  | 2 (7,7%) |  |
| II |  | 2 (7,7%) |  |
| III |  | 2 (7,7%) |  |
| IV |  | 15 (57,7%) |  |
| V |  | 4 (15,4%) |  |
| N/A |  | 0 |  |
| <u>Development of metastasis</u> |  |  |  |
| No |  | 13 (50%) |  |
| Yes |  | 13 (50%) |  |
| <u>Metastasis localization</u> |  |  |  |
| Skin (local) |  | 12 (70.6%) | 7 (31.8%) |
| Distant |  | 5 (29.4%) | 15 (68.2%) |

The variations in glycogen storage in different melanoma stages led us next to investigate whether different melanoma phenotypes exhibit distinct capacities to synthesize or store glycogen. While melanoma cells exhibit plasticity in vivo, once cultured in vitro each line maintains a fixed cell state (13) that reflects one of the phenotypes found within tumors. With this in mind, we evaluated the glycogen content *in vitro* using melanoma cell lines. Two cell lines (IGR37 and 501mel) with high levels of MITF (MITF^High^), characteristic of proliferative phenotype was compared with two MITF^Low^ melanoma cell lines (IGR39 and WM793) with invasive phenotype. The results revealed abundant glycogen stored in MITF^High^ cells and its absence in MITF^Low^ cells (Fig. 1C). Seahorse analysis of the glycolytic proton efflux rate (PER) revealed that MITF^High^ cells displayed a distinct and more glycolytic profile compared to MITF^Low^ cells (Fig. 1D). Moreover, when glucose was abundant MITF^High^ cells displayed significantly higher basal (Fig. 1E) and compensatory (Fig. 1F) glycolysis. Consistently, ATP production rate assays confirmed that the MITF^Low^ IGR39 cells are more dependent on mitochondrial OXPHOS for ATP production (Fig. S1B). The dependency of MITF^High^ cells on glycolysis as an energy source was determined measuring the extracellular acidification rate (ECAR) with a glycolytic stress test, where mitochondria are blocked. In line with our previous results, MITF^High^ cells were highly dependent on glycolysis (Fig. 1G) compared to IGR39 (MITF^Low^) that lack glycogen. MITF^High^ cells catabolize the injected glucose through the glycolytic pathway with the extrusion of protons into the surrounding medium that causes a rapid increase in ECAR. Inhibition of mitochondrial ATP production using oligomycin shifts energy production towards glycolysis, with the subsequent increase in ECAR and reveals the cellular maximum glycolytic capacity. As expected, IGR37 and 501mel cells show higher glycolytic capacity than IGR39 (Fig. 1H). The glycolytic reserve, defined as the difference between maximal glycolytic capacity and basal glycolytic rate, was dramatically reduced from MITF^High^ (IGR37 and 501mel) to MITF^Low^ (IGR39) cells (Fig. 1I), consistent with the presence of glycogen stores.

Together, the data indicate that well-defined melanoma phenotypes (proliferative and invasive) differ in their ability to store and mobilize glycogen and suggest that this metabolic adaptation could play an important role in melanoma progression.

### Proliferative, MITF^High^ melanoma cells mobilize glycogen under stress conditions

Glycogen formation begins with the critical conversion of glucose 6-phosphate (G6P) to glucose 1-phosphate (G1P), a reversible reaction catalyzed by the evolutionarily conserved enzyme Phosphoglucomutase 1 (PGM1) (Fig. 2A); subsequently, G1P is converted to UDP-glucose and used to synthesize glycogen by the enzyme Glycogen Synthase (GYS1) which elongates the chain by adding alpha 1,4 bonds. Glycogen degradation begins with the activity of the enzyme glycogen phosphorylase (PYG) which removes glucosyl units. Two isoforms PYGL, characteristic of <u>L</u>iver or PYGB, more ubiquitous and characteristic of <u>B</u>rain are expressed in melanoma cells. The G1P/G6P ratio, regulated by PGM1 activity, reflects the metabolism of glucose (predominance of glycolysis or glycogen storage) in a cell. The G1P/G6P ratio in MITF^High^ and MITF^Low^ melanoma cells was compared under conditions of glucose abundance. Coherent with previous results, a higher G1P/G6P ratio, compatible with glycogen storage or increased glycolysis was detected in MITF^High^ cells compared to MITF^Low^ (Fig. 2B). Furthermore, the levels of the key enzymes for glycogen synthesis (PGM1 and GYS1) were higher in MITF^High^ IGR37 and 501 cells than in the invasive, MITF^Low^ melanoma lines (Fig. 2C). Likewise, the levels of the glycogen degradation enzyme PYGL were increased in MITF^High^ melanoma cells (Fig. 2D). Interestingly, the invasive WM793 cells lost PYGL but increased the alternative isoform, characteristic of brain and tumor tissues, PYGB. Thus, the levels of enzymes for glycogen metabolism correlate with the capacity to store glycogen in melanoma cells. Further, RNAseq was used to compare the mRNA expression of these enzymes in our in-house panel of 12 melanoma cell lines (Fig. 2E), ranked by the expression of MITF as a marker for proliferative/differentiated cells, SOX10, a marker of the differentiated and neural crest-like states, and the AXL receptor tyrosine kinase (RTK) and SOX9 as markers of invasive phenotypes(14). Remarkably, *PGM1* and *GYS1* expression correlated with *MITF* expression in melanoma cells (Fig. 2E). Concerning the glycogen degradation enzymes, PYGL was also expressed predominantly in MITF^High^ cells, and was not found in most MITF^Low^ cell lines (Fig. 2E). Interestingly, PYGB expression appeared high in many MITF^Low^ cell lines, and low in MITF^High^ cells. An alternative panel of 53 well characterized melanoma cell lines from the Cancer Cell Line Encyclopedia (CCLE) ranked by *MITF* expression was then used to extend our findings, (Fig. 2F). The results largely recapitulated those of our in- house cell lines, with higher expression of glycogen metabolism genes in the MITF^High^ cells.

**Fig. 2.**
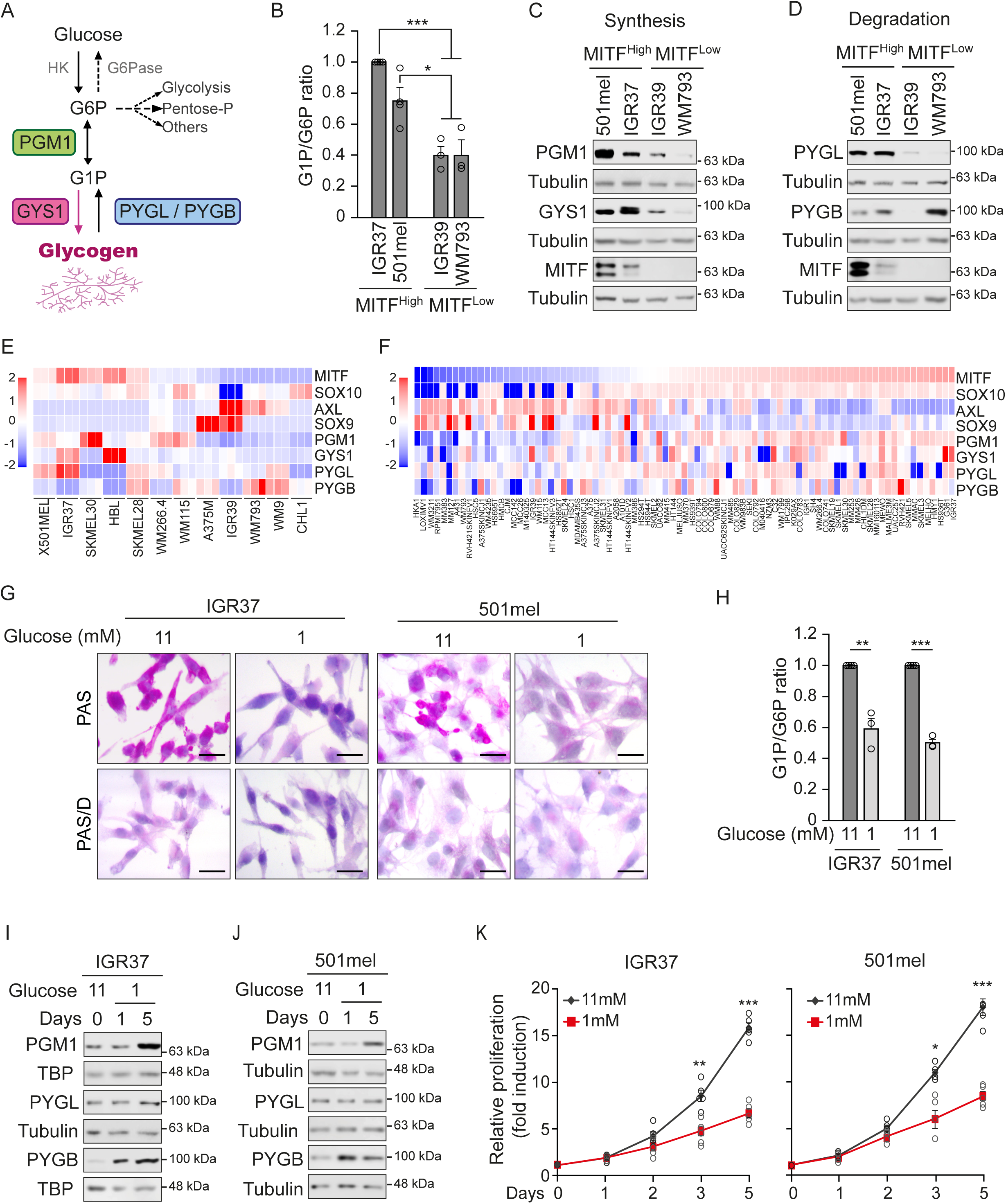
Melanoma cells express different levels of glycogen metabolism enzymes. (A) Schematic showing glycogen metabolism pathway with the main synthesis and degradation enzymes. (B) Changes in the relative amounts of Glucose 1-phosphate (G1P) and glucose 6-phosphate (G6P) in MITF^High^ (IGR37, 501mel) and MITF^Low^ (IGR39 and WM793) melanoma cells. Values represent G1P/G6P ratio, n=4 for IGR37 and 501mel and n=3 for IGR39 and WM793. (C, D) Western blot showing expression of indicated proteins in panel of IGR37 and 501mel (MITF^High^) melanoma cells. Tubulin as loading control. (E) Heatmap showing expression of MITF, PGM1, GYS1, PYGL, PYGB, AXL, SOX9 mRNA in an in-house panel of 12 melanoma cell lines measured by triplicate RNA-seq or (F) in the panel of CCLE 53 melanoma cell lines ranked by MITF expression. (G) Representative images of the variation of glycogen accumulation in MITF^High^ (IGR37, 501mel) melanoma cells in high glucose (11 mM) or low glucose (1 mM) for each cell line, staining by PAS (upper) and treated with diastase (PAS-D) (lower) to assess the specific glycogen staining. Scale bars: 25 μm. (H) G1P/G6P ratio measurement in IGR37 and 501mel melanoma cells growing in high (11 mM) or depleted of glucose (1 mM) for 24h. Values represent G1P/G6P ratio, n=3. (I, J) Western blot analysis of melanoma cells growing in high glucose (11 mM) or depleted of glucose (1 mM) for 1 or 5 days. TBP or tubulin were used as loading control. (K) Effect of glucose depletion (1 mM) on IGR37 and 501mel melanoma cells proliferation for 5 days. Black lines represent cells proliferating in high glucose (11mM). n=5. Mean ± SEM of replicates is displayed for G1P/G6P ratio (panels B and H), and for proliferation quantification (panel K). Statistical analysis by One-Way ANOVA (panel B and H) or Paired t test (panel K). *P < 0.05; **P < 0.01; ***P < 0.001.

Glycogen mobilization should provide an energetic advantage to melanoma cells under metabolic stress, facilitating tumor progression. Consequently, glycogen stores were depleted in MITF^High^ cells (IGR37 and 501mel) by culture under low glucose (Fig. 2G) with a concomitant reduction in the G1P/G6P ratio in IGR37 and 501mel cells (Fig. 2H). Under these conditions PGM1 would be needed to drive G1P from glycogen degradation towards G6P that would be able to enter glycolysis. Consistent with this, we noted increased PGM1 levels 5 days after glucose depletion (Fig. 2I-J). The key glycogen degrading enzymes appeared less sensitive to glucose depletion, although while PYGL protein levels did not vary, PYGB levels increased in cells grown in 1 mM glucose (Fig. 2I-J). Conversely, glycogen levels dramatically augmented when cells were cultured with high glucose (Fig. S2A) and the levels of the enzymes to synthesize glycogen PGM1 and GYS1 in MITF^High^ cells varied similarly (Fig. S2B). As with glucose depletion, PYG enzymes were less sensitive to glucose addition with PYGL protein levels unchanged whereas PYGB levels were reduced (Fig. S2C) as expected. By contrast, IGR39 MITF^Low^ cells were unable to accumulate glycogen despite glucose abundance (Fig. S2D) and the low levels of glycogen metabolism enzymes did not change in response to high glucose exposure (Fig. S2E-S2F). In line with this, glucose deprivation reduces the proliferation rate of MITF^High^ melanoma cells very slowly with moderate changes only appearing by 48 h (Fig. 2K).

The results indicate that MITF^High^ melanoma cells are able to accumulate glucose in the form of glycogen under nutrient rich conditions and expend it to obtain energy under glucose limiting conditions. By contrast IGR39 cells do not accumulate glycogen and do not respond to changes in glucose levels.

### The proliferative capacity of **MITF^High^** melanoma cells relies on glycogen mobilization

In melanoma and other tissues, highly proliferative cells are characterized by a metabolism that is predominantly glycolytic and strongly dependent on glucose (5). Therefore, the correlation between glycogen accumulation and greater proliferative capacity of MITF^High^ cells, suggests first that proliferative phenotype cells take up more glucose than they use for energy production or fabricating biomolecules required for cell division; and second, that the excess glucose is used to make and store glycogen that will act as a metabolic buffer by providing glucose when extracellular glucose is limiting. As such, a capacity to store glycogen will confer an advantage to proliferative cells. We therefore used Seahorse assays to analyze the metabolic plasticity of MITF^High^ cells. Glucose stress tests were used under basal conditions with or without inhibitors of glycogen metabolism. First, the glycolytic rate of IGR37 cells was determined by measuring the ECAR over time under conditions of glucose abundance (11 mM) or limitation (1 mM). The results (Fig. 3A-C) revealed that glucose depletion slightly decreased the ECAR in the short term, within the first 24 h, and at least 2 days of glucose depletion were needed for the ECAR to be abolished (Fig. 3A). Quantification of changes in ECAR revealed that long term (2 days) glucose depletion strongly reduced glycolysis by 60% (Fig. 3B) and abolished the glycolytic reserve of IGR37 cells (Fig. 3C). Similar changes with milder reductions were recorded in 501mel cells (Fig. S3A-B). In contrast, MITF^Low^ cells did not change their ECAR in response to glucose depletion in the short or long term (Fig. S3C-D). Changes in either glycolysis or compensatory glycolysis measured using the glycolytic rate assay revealed that glucose depletion for 1 day does not change the proton efflux rate (PER) (Fig S3E) or compensatory glycolysis (Fig. S3F). However, a strong reduction of 50% in both glycolysis and compensatory glycolysis was noticed after long term (2 days) glucose depletion (Fig. S3E-F). The fact that MITF^High^ cells (IGR37 and 501mel) depleted of glucose maintain their maximum glycolytic capacity and reserve for at least the first 24 h, reflects their ability to use glycogen as glucose source under stress conditions; loss of glycogen over longer periods correlates with the loss of glycolytic capacity and reserve after 2 days and with the time needed to record significant reductions in the proliferation rate (as shown in Fig 2K). Therefore, glycogen mobilization fuels the proliferation capacity of MITF^High^ cell lines and lack of glycogen correlates with low proliferation capacity in MITF^Low^ cell lines (Fig. S3G).

**Fig. 3.**
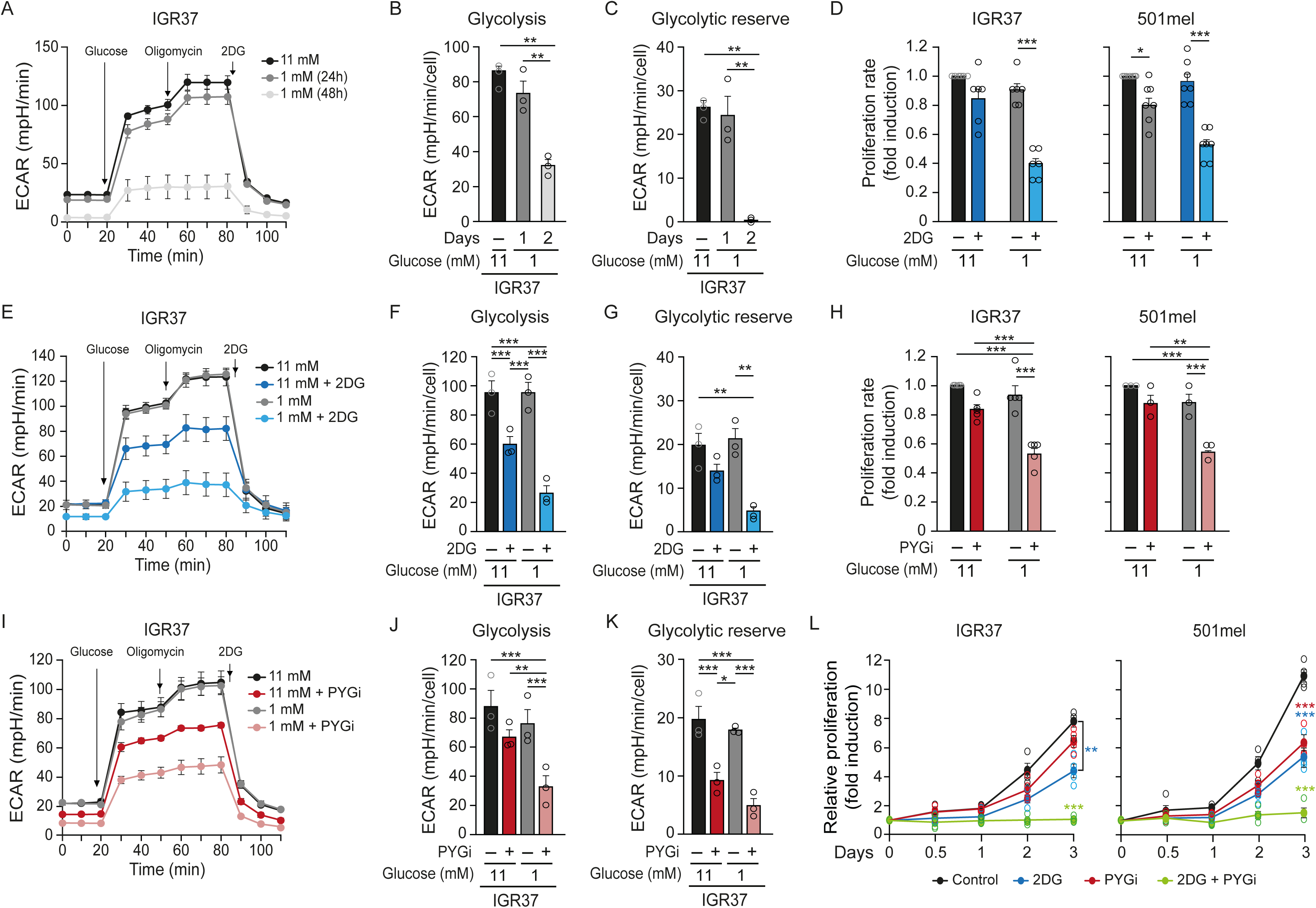
Glycogen fuels glycolysis in MITF^High^ melanoma cells to proliferate under glucose stress conditions. (A, E, I) Extracellular acidification rate (ECAR) of IGR37 melanoma cells depleted for glucose (1mM) for 1 and 2 days (A), and/or pre-treated 1.5 mM with 2-Deoxy-D-Glucose (2DG) glycolysis inhibitor (D) or 50 µM glycogen phosphorylases inhibitor (PYGi = CP-91149) for 24h (G). The media of 3 experiments is shown. (B- C, F-G, J-K) Quantification of glycolytic parameters derived from A, E or I. Bar graphs representing Basal glycolysis (B, F, J) and the glycolytic reserve (C, G, K) in each condition. n=3; error bars indicate SEM * P<0.05, **p < 0.01, ***p < 0.001. One-Way ANOVA Statistical test. (D, H) Effect of 2DG (D) or PYGi (H) on the glucose depletion induced IGR37 and 501mel melanoma cell proliferation at 24h. n=6 for IGR37 and n=7 for 501mel in D; n=5 for IGR37 and n=3 for 501mel in H; error bars indicate SEM, *p < 0.01, ***p < 0.001. One-Way ANOVA Statistical test. (L) Proliferation curves of IGR37 and 501mel melanoma cells in response to 2DG (1,5mM) (blue) or PYGi (50 µM) (red) or both (green) compared to control growing in 11mM glucose (black). n=4; error bars indicate SEM * P<0.05, **p < 0.01, ***p < 0.001. One-Way ANOVA Statistical test.

The results suggested that blocking the ability to use glycogen should alter the glycolytic profile and the proliferative capacity of MITF^High^ melanoma cells. We therefore tested this hypothesis using Seahorse analysis. Glucose depletion from 11 to 1 mM for 1 day, or 24 h treatment with 2-deoxy-D-glucose (2DG), a potent glycolysis inhibitor, did not significantly reduced the proliferation rate of IGR37 cells and decreased only mildly (less than 20%) that of 501mel cells, as expected (Fig. 3D). However, inhibition of glycolysis by 2DG in cells cultured in 1 mM glucose dramatically reduced the proliferation rate by 60% and 40% in IGR37 and 501mel respectively, unveiling the effect of 2DG in the absence of glycogen reserves. The glycolytic profile of IGR37 cells cultured with 11 mM glucose was moderately decreased upon 2DG treatment for 24 h (Fig. 3E). The ECAR during glycolysis was reduced by 30% and basal glycolysis by 40% (Fig. 3F) without significant alterations in the glycolytic reserve (Fig. 3G). However, when cells were cultured with 1 mM glucose, a strong, 75-80% reduction in ECAR was measured in glycolysis and in the glycolytic reserve was almost lost in the presence of 2DG (Fig. 3F-G).

We then more specifically targeted the ability to use glycogen and asked how this would affect the proliferation rate and glycolytic profile of MITF^High^ cells. An inhibitor specific for the enzymes that degrade glycogen (PYG) CP-91149 (15) was used to treat cells cultured as before with 11 mM or 1 mM glucose. As in the experiments with 2DG, exposure to the PYG inhibitor (PYGi) for 24 h did not change the proliferation rate of MITF^High^ (IGR37 or 501mel cells) cultured with 11 mM glucose but greatly reduced it up to 50% in cells cultured with 1 mM glucose (Fig. 3H). The glycolytic profile of IGR37 cells was modified by using CP-91149 in a similar way to 2DG, with the stronger effect in cells cultured with 1 mM glucose (Fig. 3I). Glycolysis was non significantly modified by PYGi in cells cultured with 11 mM glucose but was strongly reduced, more than 50%, in cells cultured in 1 mM glucose according to the changes in ECAR (Fig. 3J). The glycolytic reserve was also significantly reduced by 50% in cells cultured with 11 mM glucose upon treatment with CP-91149 for 24 h (Fig. 3K).

Surprisingly, combined inhibition of glycolysis and glycogen utilization using CP-91149 and 2-DG simultaneously, completely blocked the proliferative capacity of MITF^High^ IGR37 and 501mel cells cultured with 11 mM glucose (Fig. 3L), highlighting the importance of glycogen usage as an energy source under energetic stress.

Thus, blockade of glycogen metabolism in MITF^High^ melanoma cells when cultured under glucose stress (1 mM) strongly reduces the glycolytic capacity by 24 h, with a later reduction in the proliferation rate. The results indicate that glycogen fuels glycolysis when the availability of extracellular glucose becomes limiting as would occur during tumor expansion.

### Loss of capacity to use glycogen induces a phenotype switch towards invasiveness in melanoma cells

So far, the results indicate that glucose stress conditions regulate the key enzymes for glycogen metabolism in MITF^High^ melanoma cells to enable them to use glycogen stores to support proliferation. Whether the expression of these enzymes is upregulated in proliferative tumors is unknown. We therefore interrogated the TCGA skin cutaneous melanoma cohort (SKCM) that revealed a strong positive correlation between the Verfaillie proliferative signature gene set (16) and *GYS1* mRNA expression (r=0.380; p< 2.2x10^-16^), (Fig. 4A). Likewise, the Verfaillie proliferative signature also positively correlated with PYGB mRNA expression (r=0.236; p=2.05 x 10^-7^) whereas PYGL rendered a weak correlation. Importantly, a positive correlation between GYS1 and MITF (r=0.250; p=3.3 x 10^-8^) and although weaker, also between PYGB and MITF, (r=0.126 and p=0.006) highlighted the link between glycogen metabolism and proliferative melanomas (Fig. S4A-B).

**Fig. 4.**
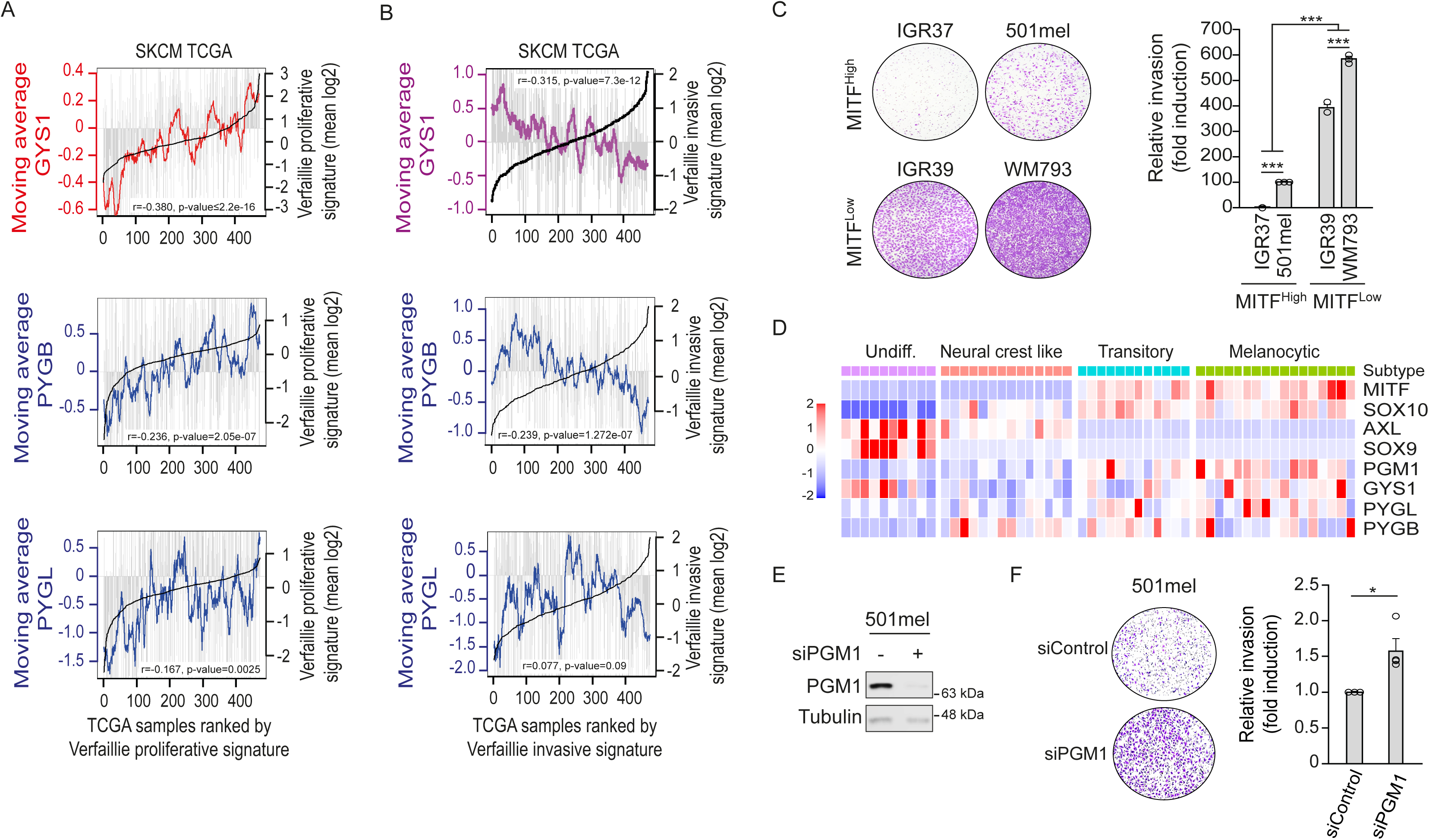
Loss of the ability to use glycogen as energy source induces a phenotype switch in melanoma cells. (A- B) Plot showing GYS1, PYGL or PYGB mRNA expression (grey bars) in the TCGA melanoma cohort ranked by Verfaillie proliferative (A) or invasive (B) signature expression (black line). The moving average of each per 20 melanoma window is indicated by the coloured lines. P value and correlation coefficient (rho) are indicated. (C) Matrigel transwell invasion assays showing invading cells stained with crystal violet. Representative images of the basal invasion of melanoma cells growing in 11mM glucose for 24 hours is shown (left). Quantification from n=3 biological replicates except in the right.; error bars = SEM; * =P< 0.5, ** =P< 0.1, *** = p< 0.01. One-Way ANOVA Statistical test. (D) Heatmap showing relative expression of GYS1, PGM1, PYGB, and PYGL in the Tsoi et al (17) cell lines using MITF, SOX10, SOX9 and AXL expression as markers of indicated phenotypes. (E) Western blot showing PGM1 protein levels in 501mel melanoma cells transfected control or PGM1 targeting siRNAs. Tubulin as loading control. (F) Matrigel transwell invasion assays showing invading cells stained with crystal violet. Cells were transfected with control or PGM1targeting siRNAs. n=3; error bars indicate SEM * P<0.05, **p < 0.01, ***p < 0.001. Paired t test.

Consistent with these observations, the Verfaillie invasive gene expression signature significantly and inversely correlated with GYS1 (r=0.315; p=7.3 x 10^-12^) and PYGB (r=0.239; p=1.27 x 10^-7^) but not with PYGL, (Fig. 4B). The data imply that the importance of glycogen metabolism is reduced or absent in de- differentiated, invasive, and slow cycling cells. MITF^Low^ cells have a higher invasive capacity than MITF^High^ cells (Fig. 4C) which is consistent with their absence of glycogen stores and the low levels of key enzymes for glycogen metabolism.

The possible association of glycogen metabolism enzymes with defined melanoma cell phenotypes was explored in a panel of 53 well characterized melanoma cell lines subdivided into 4 different phenotypes (17) with *MITF*, *SOX10*, *SOX9* and *AXL* used as markers of cell state (Fig. 4D). The results indicate that PYG enzymes for glycogen degradation are present in differentiated melanocytic phenotypes, which showed high PYGL and low PYGB mRNA levels; PYGL tended to decrease and PYGB to increase in transitory and neural crest phenotypes and both PYGL and PYGB were absent in undifferentiated phenotypes. For glycogen synthesis, GYS1 mRNA levels were also high in melanocytic phenotypes. The presence of high levels of GYS1 mRNA in undifferentiated phenotypes that lack PGM1 (to provide substrate for GYS1 reactions) is interesting and puzzling. Remarkably, PGM1 mRNA expression, like PYG expression, was largely restricted to MITF^High^ melanocytic and transitory melanoma cell phenotypes that lack expression of AXL, a hallmark of melanoma invasion and therapy resistance (Fig. 4D) (14,18). This prompted us to ask asked whether PGM1 depletion might impact melanoma cell phenotype. Knockdown of PGM1 in MITF^High^ melanoma cells (Fig. 4E) before challenging them with Matrigel invasion assays, revealed more than 50% increased invasiveness in cells depleted of PGM1 (Fig. 4F) despite the abundance of glucose (11 mM).

The results indicate that loss of PGM1 in MITF^High^ cells favor a phenotypic switch towards an invasive phenotype.

### PGM1- and PYGB-dependent glycogen mobilization during human melanoma progression has prognostic value

In many cancer types other than melanoma, high tumor expression of enzymes related to glycogen metabolism is associated with worse patient outcomes (19), but the underlying mechanisms remain unclear. To understand whether and how the gene expression of enzymes required for glycogen synthesis (*GYS1*, *PGM1*) or degradation (*PYGB*, *PYGL*) is associated with patient outcome we explored the TCGA database with 571 human cutaneous melanoma samples matched with non-cancerous tissue. TNM plot analysis revealed mRNA increases from normal skin to tumor tissue for *PGM1* (Fig. 5A), *GYS1* (Fig. 5B), *PYGB* (Fig. 5C) and *PYGL* (Fig. 5D) that were significant in all cases (p<0.001); despite the under representation of melanocytes in normal skin, the data implies that expression of genes for glycogen metabolism is important in primary tumors. Comparison between primary tumors and metastases revealed significant (P<0.05) changes in the expression of enzymes involved in glycogen synthesis, with increased *PGM1* (Fig. 5A) and decreased *GYS1* (Fig. 5B) expression, whereas no significant changes were observed in the expression of genes encoding glycogen degrading enzymes *PYGB* or *PYGL* (Fig. 5C-D). The data is compatible with an increased need to express enzymes for glycogen metabolism early in tumor formation (from non-malignant to tumor) and the progressive loss of glycogen metabolism towards metastasis. The tendency to decrease and the ample variation in metastasis for gene expression of glycogen metabolism enzymes is indicative of a heterogeneity of adaptive metabolic responses that may vary with the new niche.

**Fig. 5.**
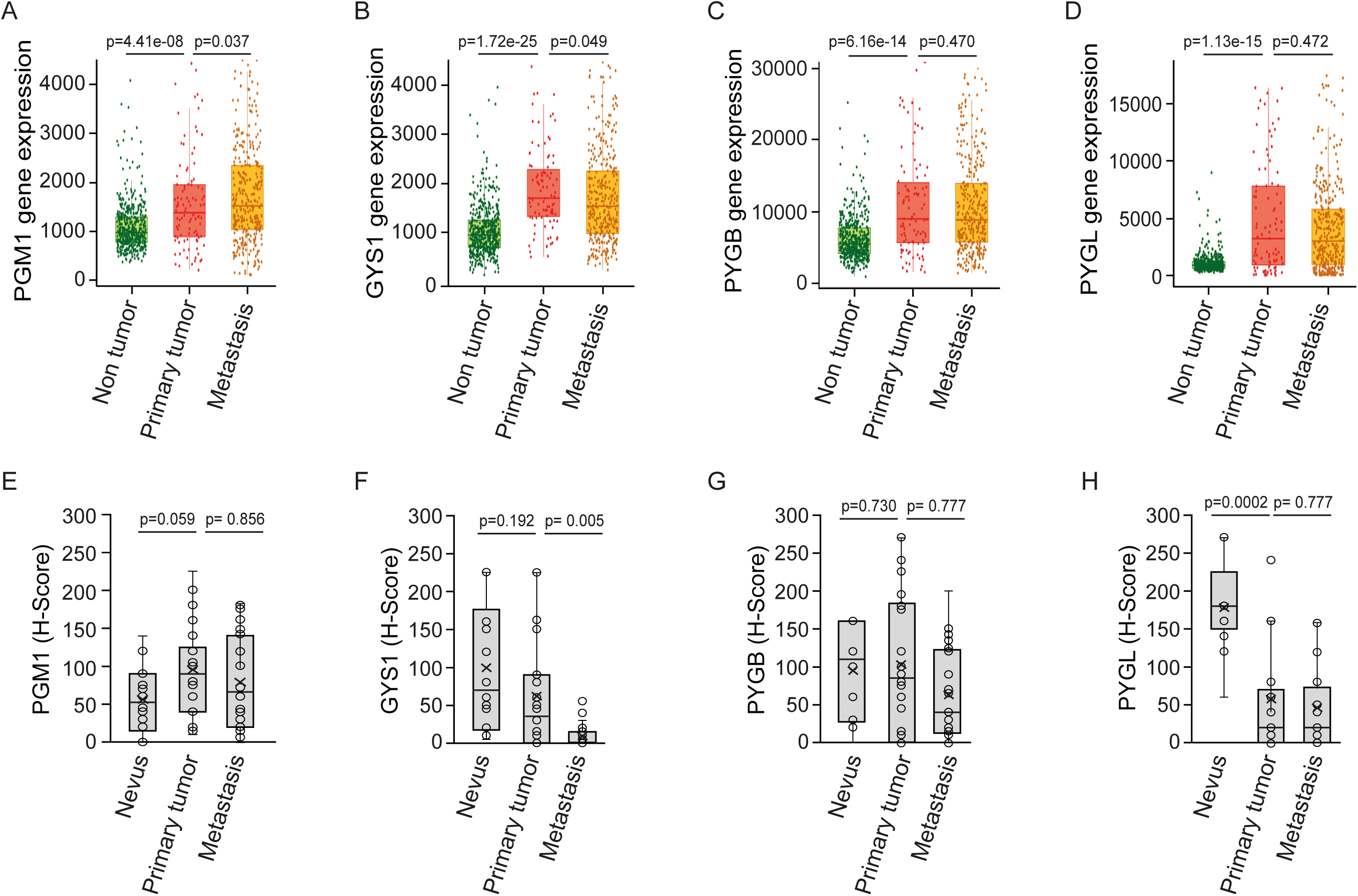
Glycogen metabolism enzymes levels in human melanoma samples. (A-D) TNM plot analysis of PGM1 (A), GYS1 (B), PYGB (C) or PYGL (D) gene expression in 377 samples of non-cancer tissue (normal) versus 211 melanoma tissue (tumor) and 111 metastatic tissues (metastatic). Data was collected from TCGA database. Expression is shown as transcripts per million (TPM) in log2. Samples were compared with Mann-Whitney U test. The statistical significance cut off was set at p < 0.01. (E-H) Profile for PGM1 (E), GYS1 (F), PYGB (G) and PYGL (H) content from nevus, melanoma and metastatic human samples as in 1B. The profile was obtained from Histoscore (H-Score) variations. Mann-Whitney U statistical test. The statistical significance cut off was set at p < 0.01.

The multiplicity of mechanisms that regulate enzymes leads to frequent discrepancies between mRNA and protein levels. We therefore compared by immunohistochemical analysis pre-malignant human nevi (n=14), to primary tumors (n=27) and metastatic samples (n=28) in a tissue microarray (TMAs) from the Biobank of Hospital Clínico San Carlos (Table 1). Comparison of nevi with primary melanoma samples showed increased PGM1 enzyme levels with a high trend towards significance (Fig. 5E) in line with the mRNA expression data, although at the step between the primary melanoma and metastasis, protein levels tend to decrease (non- significantly) whereas mRNA expression was increased. GYS1 protein levels (Fig. 5F) showed a tendency to decrease (although non-significant, probably due to the reduced number of samples), but which contrasted with the significant increase recorded for mRNA expression in Fig. 5B. A tendency that is supported by a strong and significant (p=0.005) decrease between primary melanoma and metastases, which suggests that metastasis does not require glycogen synthesis or that glycogen suppresses metastatic dissemination.

For the glycogen degrading enzymes, PYGB did not show significant differences between nevi and primary melanoma or metastases (Fig. 5G). However, PYGL (Fig. 5H) revealed a very significant (p=0.0002) decrease in protein levels from nevi to primary melanoma which contrasted with the increased mRNA expression presented in Fig. 5D. The step from tumor towards metastases was characterized by a tendency to decrease (non-significant) PYGL mRNA expression and protein levels, suggesting again that glycogen metabolism is non-essential for metastases.

Interestingly, the levels of PGM1 and PYGB proteins showed a very significant positive correlation (r=0.578, p<0.0001) (Fig. S5A). Note the dual role of PGM1, which may participate in glycogen synthesis by converting G6P to G1P, or in glycogen degradation by converting G1P into G6P.

Globally, the data suggested that the capacity to store and use glycogen is important in the early stages of melanoma, where MITF^High^ melanoma cells use glycogen as an energy source to proliferate, and this capacity loses importance as the disease progresses. To explore this, we evaluated the possible association between the levels of the enzymes (GYS1, PGM1, and PYGB) and clinicopathological features of disease progression including the invasiveness through the skin (measured by Clark levels), establishment of metastases and distance of metastasis. GYS1 levels did not associate with any of these previous features (Fig. S5B). However, both PGM1 and PYGB were positively and significantly associated with Clark level, (Fig. 6A-B) when melanoma primary tumors were analyzed separately. Importantly, this correlated with the loss of PAS staining (Fig. S5C), featuring the importance of glycogen usage for the progression of melanomas. Furthermore, both PGM1 and PYGB positively correlated with increased capacity to establish metastases (Fig. 6C). The fact that primary tumors that develop metastasis exhibit higher levels of PGM1 and PYGB is consistent with an advantage of tumors that metabolize glycogen to progress. Higher PGM1 and PYGB levels correlate specifically with primary tumors that develop local metastases (Fig. 6D). In contrast, melanomas that developed distant metastasis in lung, liver or brain, which are more aggressive and drive worse patient outcomes, exhibited lower PGM1 and PYGB levels (Fig. 6D), which is consistent with our data that loss of PGM1 favors invasiveness (Fig 4F).

**Fig. 6.**
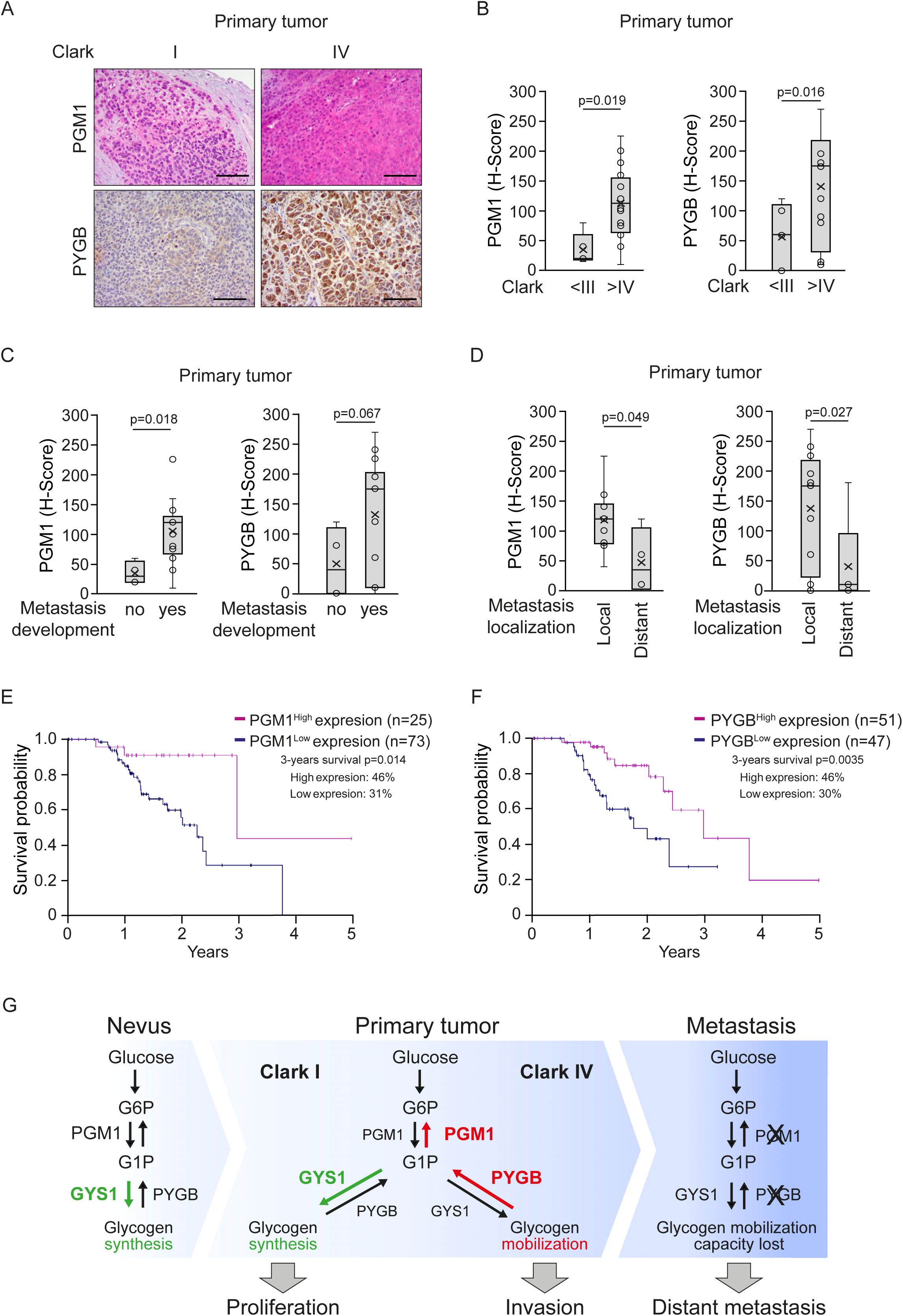
Clinical relevance of PYGB and PGM1 levels in human melanoma samples. (A) Representative micrographs from immunostaining for PGM1 or PYGB on human samples of melanoma with I or IV Clark level. Counterstaining with hematoxylin stains nuclei in dark blue and cytoplasm in light blue. Scale bars: 50 μm. (B-D) Profile for PGM1 or PYGB content from melanoma primary tumor samples with low or high Clark level (B), that developed metastasis or not (C) or the metastasis localization (D). Statistical analysis by Man Whitney U-test; *p<0.05, **p<0.01 and ***p<0.001. (E-F) Kaplan-Meier curves displaying the estimated survival probability for melanoma patients performed using the best cut-off point and data from 102 melanoma patients from the The Human Protein Atlas; PGM1^High^ (n = 25); PGM1^Low^ (n = 73) in E; PYGB^High^ (n = 51); PYGB^Low^ (n = 47) in F. (G) Glycogen metabolism changes during melanoma progression. Nevus and primary melanomas (Clark I) exhibit high glycogen accumulation, correlating with higher GYS1 levels in proliferative stages. Primary tumors that go on to develop metastases exhibit higher levels of PGM1 and PYGB than those which do not develop metastases; both PGM1 and PYGB are required for glycogen mobilization leading to glycogen depletion and associated loss of metabolic buffering that suppresses invasion. Furthermore, melanomas that develop distant metastases (more aggressive) and their metastases (from lung, brain or liver) exhibit lower levels of PGM1 and PYGB, which associates with worse prognosis. Green arrows indicate glycogen synthesis, red lines indicate glycogen degradation. Glycogen synthesis enzymes are highlighted in green and for degradation in red.

Analysis of metastases samples rendered similar results. Skin metastases (less aggressive and with a better prognosis) exhibited higher PGM1 and PYGB levels than distant metastases from ganglion, liver or lung (Fig. S5D). The results highlight the likely importance of the capacity to degrade glycogen for early steps of tumorigenesis. Survival data obtained through Kaplan-Meier analysis revealed that patients with high levels of PGM1 and PYGB have increased survival over those with low levels (Fig. 6E-F).

Taken together, the results indicate that glycogen metabolism is critical for cancer cell proliferation and suppresses invasiveness at early stages; loss of glycogen stores, or an inability to mobilize glycogen, facilitates a phenotype switch from proliferation to invasion (Figure 6G). The data also reveal a prognostic value for evaluation of glycogen stores in melanoma using simple PAS staining as well as immunohistochemistry for PGM1 and PYGB.

## Discussion

This work demonstrates that in melanoma, proliferative (MITF^High^) phenotype cells are especially dependent on glucose metabolism and store glucose as glycogen. Glycogen provides a survival advantage under nutrient limiting conditions such as glucose depletion that would be expected during tumor progression. Since glucose is the most important fuel and carbon source for cellular proliferation, the availability of an alternative immediately usable glucose source such as glycogen should be highly beneficial for proliferative cancer cells. This raises the possibility that any switch to an invasive phenotype in response to glucose limitation could be limited or delayed if glycogen were available. Consistent with this, we show that in melanoma loss of the capacity to mobilize glycogen favors invasiveness and associates with worse prognosis.

In support of the idea that glycogen breakdown is important for tumor growth in primary tumors, our results indicate that glycogen content decreases in advanced primary melanomas (Clark IV) and metastases compared to nevi or early melanoma stages (Clark I), consistent also with the idea that glycogen suppresses invasiveness. This is in line with the reported correlation between glycogen accumulation and improved survival in other cancer cells in response to nutrient stress (20,21) or as an adaptation to a new metastatic niche (22). Disappearance of glycogen stores during malignant transformation was previously reported in epithelial intestinal cells (23) and now we report that in melanoma, invasive capacity inversely correlates with the presence of glycogen stores both in vitro and ex vivo. The data suggests that glycogen might suppress invasiveness in melanoma or that the loss of the ability to mobilize glycogen promotes an invasive phenotype.

Glucose depletion during melanoma progression requires adjustments in the net activity of critical glycogen metabolism enzymes such as PGM1, PYGs and GYS. PGM1 catalyzes the interconversion of G1P and G6P to regulate an important glucose trafficking point: feeding glucose into glycogen synthesis under extracellular glucose replete conditions, or conducting glucose away from glycogen degradation towards glycolysis and biosynthetic pathways on glucose limitation (24). Hence, proliferative, MITF^High^ melanoma cells exhibit increased PGM1 protein levels and elevated glycogen turnover. This allows glucose buffering for cell proliferation and suggests a pro-survival role for PGM1 in melanoma upon glucose deprivation, consistent with other tumors (20,25,26). Our data indicate that PGM1 levels are elevated in primary melanomas relative to metastases. In line with that, bioinformatic analysis of melanomas showed that *PGM1* expression is upregulated in well-defined proliferative phenotypes and lost in more invasive phenotypes. The fact that in MITF^High^ proliferative melanoma cells, PGM1 knockdown drove increased invasiveness highlights the importance of losing glycogen mobilization capacity for a phenotype switch towards invasion and defines a lack of PGM1 activity as a critical driver for disease progression. Coherent with these results Kaplan Meier analysis of the melanoma collection in the Human Protein Atlas reveals that survival of melanoma patients strongly correlates with a high content of PGM1, whereas patients with low PGM1 expression have significantly lower survival. The results are likely to impact in diagnosis. Also, the identification of PGM1 and the interconversion G6P/G1P as critical for invasiveness, may offer new therapeutic targets for melanoma.

Glycogen phosphorylases (PYG) catalyze glycogen degradation to control the exhaustion of glycogen stores Two isoforms, PYGL and PYGB are present in melanoma and differ in their allosteric regulation. PYGL activity is hormonally controlled by insulin and glucagon signaling, while PYGB activity is dependent on intracellular nutrient availability (27), through the ratio of AMP:ATP. This mode of allosteric regulation confers an advantage to PYGB in cancer cells with a concomitant shunt in glycogen catabolism. Therefore, PYGB upregulation may be crucial to sustain proliferation, that will deplete glycogen ultimately leading to invasion (22,28). In line with the differential regulation of PYG isoforms, our results indicate that while PYGL levels are maintained or downregulated in melanoma, PYGB levels are upregulated, which is consistent with increased PYGB levels reported in other cancers (22,29,30), although the mechanism underpinning cancer progression by PYGB is unclear. To further our understanding on the role of PYGB in cancer progression, our results demonstrate that PYGB is increased specifically in proliferative phenotypes where its upregulation in response to decreased glucose availability helps sustain proliferation in melanoma. The results are further supported by the fact that PYGB and PGM1 levels vary in parallel, as if equally required for survival of this phenotype. Our results indicate that PYGB increases during transformation from nevus to primary melanoma and disappears during melanoma progression to distant metastases in parallel to PGM1. On the other hand, PYGL levels are considerably decreased already in the early stages of transformation, suggesting a reduced importance in cancer cell survival. In line with our results, Kaplan-Meier analysis indicates that high levels of PYGB strongly predict better survival in melanoma patients. However, other authors reported an involvement of PYGB in cancer progression and metastasis (22,31–33) and associated increased PYGB levels to worse prognosis for other tumors (34). Our model of melanoma is specially well suited for these studies because, unlike many other cancer models, cancer cell phenotypes and phenotype-switching are very well-defined.

In view of our data suggesting that glycogen stores might suppress invasiveness, the potential of glycogen catabolism for cancer biology should be thoroughly explored. Glycogen phosphorylase inhibitors CP-91149 and CP-320626 have shown potent antiproliferative effects in non-small lung carcinoma, pancreatic cancer and HCC (35–37). However, in melanoma, CP-91149 only decreased cell proliferation under glucose-limiting conditions in proliferative MITF^High^ cells. The data indicated that proliferation was fed by external glucose undergoing glycolysis and revealed the importance of glycogen stores under energetic stress. 2-DG is used in preclinical studies to significantly inhibit glycolysis and ATP synthesis (38), however in melanoma MITF^High^ cells, 2-DG did not have major effects on proliferation, likely due to the glucose supply provided by glycogen stores. This may explain contradictory results obtained in clinical trials (39,40). Interestingly, combined inhibition of glycolysis and glycogen utilization, by using CP-91149 and 2-DG simultaneously, completely blocked the proliferative capacity of proliferative MITF^High^ melanoma cells. This indicates that PYGB is not essential for the survival of melanoma cells, but it is important in early stages for proliferation. In support, a highly significant correlation between PGM1 and PYGB suggests their co regulation as part of a common regulatory axis.

Although CP-91149 has been explored as an adjuvant agent in kidney and ovarian (41) cancer, and has shown the potential to restore drug sensitivity to sunitinib in resistant ccRCC cell lines in vitro (42), or to synergize with sorafenib in HepG2 cells (43), only one glycogen phosphorylase inhibitor, flavopiridol, has been tested in clinical trials for prostate cancer, renal cell carcinoma and colorectal cancer (44–46). However, since flavopiridol is also a cyclin-dependent kinase inhibitor (47), it is unclear how it exerts its antitumorigenic activity. Nevertheless, in clinical trials, treatment with flavopiridol in combination with other drugs was safe and showed some efficacy in urological cancers (44–46). Its potential should therefore be further explored for cancers with overactive glycogen metabolism. However, some important problems must be solved first, including directing the inhibitor only to the tumor and avoiding compensatory metabolic pathways. This is important because glycogen metabolism is needed to maintain glucose homeostasis and its inhibition may cause life threatening hypoglycemia with compensatory metabolic pathways appearing to favor invasiveness. Overall, it is plausible that PYG inhibitors should be considered as a potential adjuvant agent improving the efficacy of standard chemotherapeutics.

Taken together, decreased glycogen stores were directly associated with invasiveness in human primary melanomas evaluated by Clark level. Glycogen depletion correlated with increased PGM1 and PYGB in primary melanoma tumors that progressed to metastasis, highlighting the likely importance of glycogen usage for early stages of melanoma progression. Furthermore, the higher the levels of PGM1 and PYGB the higher the probability that the primary tumor would develop distant metastases and worse patient outcome. Low PGM1 or PYGB expression associates with worse overall survival of melanoma patients, consistent with loss of metabolic buffering capacity in late melanoma stages. Therefore, we identify here PGM1 and PYGB levels as key sensors that balance proliferation with carbon availability in melanoma cells, and glycogen metabolism (storage or breakdown) as a critical hub that sustains survival and growth and which suppresses invasion. Metabolic buffering via the accumulation of glucose stores in the form of glycogen therefore suppress the ability of melanoma cells to undergo a proliferative to invasive phenotypic transition.

Cancer progression depends on metabolic reprogramming. Inside the tumor, proliferative cells will engender alternative phenotypic states to become invasive and/or resistant to therapies (4). The ability of cancer cells to store vital metabolites in forms that can be readily converted to glucose and glutamine for the later use represents a great advantage. A metabolic buffer will enable cells to delay the decision to enter an invasive state, which is associated with many potentially lethal stresses including anoikis, ferroptosis or an unresolved integrated stress response (4,5). As revealed here, metabolic buffering is likely to play a key role in determining the timing of a proliferative to invasive phenotype switch in a range of cancers beyond melanoma. As such, inhibitors of glycogen degradation may improve the success of current anti-cancer treatments. Our results therefore have major implications for our understanding of cancer progression and therapy resistance.

## Material and methods

### KEY RESOURCES TABLE

KEY RESOURCES TABLE:
| REAGENT or RESOURCE | SOURCE | IDENTIFIER | RRID |
| --- | --- | --- | --- |
| **Antibodies** |  |  |  |
| Rabbit monoclonal anti-GYS1 | Cell Signalling | Cat# 3886 | AB_2116392 |
| Rabbit monoclonal anti-Lamin B1 | Invitrogen | Cat# 702972 | AB_2784553 |
| Mouse monoclonal anti-MITF | Abcam | Ab12039 | AB_298801 |
| Rabbit monoclonal anti-PGM1 | Abcam | Ab192876 | AB_2893201 |
| Rabbit polyclonal anti-PYGB | Sigma-Aldrich | HPA031067 | AB_2673722 |
| Rabbit polyclonal anti-PYGL | Sigma-Aldrich | HPA000962 | AB_1079722 |
| Rabbit Polyclonal TFIID-TBP (N12) | Cell Signaling | Cat#44059 | AB_2799258 |
| Mouse Monoclonal anti-α-Tubulin | Sigma | Cat# T5168 | AB_477579 |
| Goat Anti-Mouse IgG (H+L) HRPO | BIO-RAD | Cat #170-6516 | AB_11125547 |
| Goat Anti-Rabbit IgG (H+L) HRPO | BIO-RAD | Cat #170-6515 | AB_11125142 |
| **Chemicals, Peptides, and Recombinant Proteins** |  |  |  |
| Antimycin A | Sigma | Cat #A8674 |  |
| 2-Deoxy-D-Glucose | Sigma | Cat #D8375 |  |
| Bradford | Sigma | Cat#B6916 |  |
| Corning Matrigel Invasion Chamber | Corning | Cat #<br>45354480 |  |
| Crystal violet | Sigma | Cat # C6158 |  |
| D – (+) - Glucose | Sigma | Cat #G8769 |  |
| Diastase | Sigma | Cat #09962 |  |
| Fetal Bovine Serum | Sigma | Cat# F7524 |  |
| Formalin | Sigma | Cat #HT501128 |  |
| Glycogen Phosphorylase inhibitor (CP-91149) | Selleckhem | Cat #S2717 |  |
| Hematoxylin | Sigma | Cat #GHS316 |  |
| Harris Hematoxylin | Sigma | Cat #HHS32 |  |
| HRP Magenta chromogen | Dako Agilent | Cat #GV900 |  |
| JetPrime PolyPlus reagent | Genycell Biotech | Cat# 114-15 |  |
| L-Glutamine (200 mM) | LONZA | Cat# BE17-605E |  |
| MEM NEAA | GIBCO | Cat# 11140-035 |  |
| Oligomycin A | Sigma | Cat #75351 |  |
| PBS | Corning | BE02-017F |  |
| Penicillin-Streptomycin | LONZA | Cat# DE17-602E |  |
| Periodic acid | Sigma | Cat #3951 |  |
| Peroxidase blocking reagent | Dako Agilent | Cat #S202386 |  |
| Protease inhibitor Cocktail | Roche | Cat#04693132001 |  |
| Rotenone | Sigma | Cat #R8875 |  |
| RPMI 1640 media | Corning | Cat# 5510-041-cv |  |
| Shift Reagent | Sigma | Cat #3952016 |  |
| Tween 20 | Sigma | Cat #P1379 |  |
| **Critical Commercial Assays** |  |  |  |
| Clarity ECL detection kit | BIO-RAD | Cat#170-5060 |  |
| Glucose-1-Phosphate (G1P) Colorimetric Assay Kit | Sigma | Cat# MAK098 |  |
| Glucose-6-Phosphate (G6P) Colorimetric Assay Kit | Sigma | Cat# MAK014 |  |
| EnVision + Dual link System HRP | EnVision, Dako,<br>Agilent | Cat#K4065 |  |
| **Cell lines (*Homo-sapiens*)** |  |  |  |
| Pierce BCA Protein Assay Kit | ThermoFisher | Cat#23225 |  |
| 501mel (Female). Human melanoma cell line | Obtained from<br>Ruth Halaban,<br>Yale | (48) | CVCL_4633 |
| IGR37 (Male). Human melanoma cell line. | Obtained from<br>Luisa<br>Lanfranccone | (49) | CVCL_2075 |
| IGR39 (Male). Human melanoma cell line. | Obtained from<br>Luisa<br>Lanfranccone | (49) | CVCL_2076 |
| WM793 (Male). Human melanoma cell line | Obtained from<br>Meenhard Herlyn | (50) | CVCL_8787 |
| **Oligonucleotides** |  |  |  |
| ON-TARGETplus Non-targeting Pool (negative control siRNA) | Dharmacon Inc | D-001810-10-05 |  |
| SMARTpool: ON-TARGETplus human PGM1 (5236) siRNA | Dharmacon Inc | L-010925-01-0005 |  |
| **Software and Algorithms** |  |  |  |
| Agilent Seahorse Analytics | Agilent | Seahorseanalytics.agilent |  |
| Gene Cluster 3.0 | deHoon et al.,<br>National<br>Institutes of<br>Health, 2004. |  |  |
| GraphPad Prism software | GraphPad<br>Software | https://www.graphpad.com | SCR_002798 |
| ImageJ software | N/A | https://imagej.nih.gov/ij/download.html | SCR_003070 |
| ImageLab | Bio-Rad | ChemiDoc XRS+ System | SCR_014210 |
| LAS AF software | Leica | SP5 | SCR_013673 |
| SPSS Statistics version 26 | IBM | N/A |  |
| Molecular Signatures | mSigDB | http://www.broadinstitute.org/msigdb |  |
| RNAseq data | cBioportal | http://www.cbioportal.org | SCR_014555 |
| **Other** |  |  |  |
| TCGA analysis R-script | N/A |  |  |

### Cell lines

All human melanoma cell lines were cultured in RPMI supplemented with 10% fetal bovine serum (FBS) plus 1% penicillin–streptomycin and maintained in 5% CO2 environment at 37°C. All cell lines were authenticated by STR profiling and mycoplasma contamination tests by PCR were run monthly to ensure no contamination.

### RNAi gene silencing

Cells plated in six well plates at 50% confluence were transfected with siRNA using JetPRIME Polyplus reagent (Genycell Biotech, Santa Fe, Granada, Spain), following the manufacture instructions. After 2 days cells were treated and collected to perform invasion assay or to be analyzed by western blot. Specific siRNA oligonucleotides for PGM1 and control siRNA were obtained from Dharmacon Inc.

### Preparation of cell extracts

Cells were washed with iced PBS before extract preparation and scraped in RIPA buffer (10 mM Tris-HCl pH 7.4, 5 mM EDTA, 5 mM EGTA, 1% Triton X100, 10 mM Na4P2O7, pH 7.4, 10 mM NaF, 130 mM NaCl, 0.1% SDS, 0,5% Na-deoxycholate). After 5 min on ice, cells were pelleted (12,000 rpm for 5 min, 4°C) and the supernatant was directly used as whole cell extract or frozen at -80 °C.

### Western blot

Protein lysates were subjected to 10 to 15% polyacrylamide SDS-PAGE. Proteins were transferred onto polyvinylidene difluoride membranes. Membranes were blocked with 5% non-fat milk in T-TBS containing 0.1% Tween 20 and probed with the appropriate primary antibodies (see Key Resources Table) overnight at 4°C. The specific bands were analyzed using ChemiDoc Imaging Systems (Bio-Rad).

### Crystal violet staining and cell density quantification

30,000 cells were plated in a 12-well plate for 24h, then depleted from glucose and/or treated with CP-91149 or 2-DG over 5 days as indicated. The medium was replaced every 3 days. Plates were collected at time 0, 12, 24, 48, 72, or 120 h, fixed with 4% PFA, stained with 0.1% crystal violet for 15-30 min, washed with PBD and dried. Crystal violet was resuspended methanol, transferred to p96 plates and analyzed by a Spectra FLUOR (Tecan) at 570 nm. Viability was measured by duplicate.

### Matrigel invasion assays

Matrigel invasion assays were performed using invasion chambers from BD Biocoat 8 µm membrane inserts with Matrigel coating. Melanoma cells treated or transfected as indicated were cultured in medium without 0.2% FBS overnight and 100.000 cells were seeded per insert. Medium with 10% FBS was added to the bottom of the inserts as chemoattractant. After 22 h, chambers were fixed in 4% paraformaldehyde for 2 min, washed in PBS, permeabilized 20 min with 100% methanol, washed again, stained with 1% crystal violet for 20 min and washed in PBS. Cells remaining above the insert were removed by gentle scraping with a sterile cotton swab. Slides were mounted and images were acquired using a Zeiss optical microscope with a 5x objective. Cells were manually counted using ImageJ software. At least three biological replicates were performed.

### G6P and G1P measurement

Glucose 1 phosphate and glucose 6 phosphate cells concentrations were determined by colorimetric assays following the manufacturer instructions (Sigma-Aldrich, St. Louis, MO, United States). Cells growing in 11 mm glucose were depleted for glucose (1mM) for 24 hours. 1x10^6^ cells were homogenized in 2–3 volumes of ice-cold PBS and centrifuge the samples at 13,000 × g for 10 minutes to remove insoluble material. 25 µl of sample were diluted with 25 µl of assay buffer and incubated with 50 µl of the appropriate Reaction Mix, incubate the reaction for 30 minutes at room temperature and measured the absorbance at 450 nm. Glucose-6- Phosphate and Glucose-1-Phosphate standards were used to plot a standard curve. Measurements were determined in duplicates in at least three different experiments.

### PAS and PAS-D staining

Melanoma cells seeded in coverslips were washed 3 times, fixed with 4% paraformaldehyde in PBS (pH 7.4) for 10 min, and washed again. For PAS-D staining, slides were incubated with 0.2% α-amylase (Diastase, Sigma-Aldrich) in PBS for 30 min at 37°C and rinsed in tap water and then with distilled water. For PAS staining, this step was performed with PBS. Slides were then oxidized for 5 min with 1% solution of periodic acid (Sigma-Aldrich), washed with tap water for 3 min, washed with distilled water for 1 min, and incubated in Schiff’s reagent (Sigma-Aldrich) for 20 min. Slides were washed first with distilled water for 5 s and then with tap water for 10 min. Counterstaining was performed with Hematoxylin Solution (Sigma-Aldrich) for 2 min and washed 3 times for 5 min with PBS.

For tissue sections, slides were deparaffinized and hydrated before diastase digestion (2% α-amylase 1h at 37°C) and PAS staining: 5 min in 1% solution of periodic acid followed by washes and incubation in Schiff’s reagent for 10 min. Slides were counterstained with Hematoxylin for 30 s. Images were acquired using a microscope (Zeiss) with a 40× objective for cultured cells or 20× for tissue sections.

### Seahorse experiments

Cellular metabolism was analyzed by measurement of glycolytic rate for basal conditions and compensatory glycolysis following mitochondrial inhibition (Agilent Seahorse XF Glycolytic Rate Assay), ATP production from mitochondrial oxidative phosphorylation (OXPHOS) and glycolysis (Agilent Seahorse XF Real-Time ATP Rate Assay) and key parameters of glycolytic function (Agilent Seahorse XF Glycolysis Stress Test). Depending on the cell line and assay type, 15,000 to 25,000 cells were seeded in 100 µL of RPMI per well of 24-well plates (XF24) and after 4-6 hours of incubation at 37°C, once the cells were adhered to the plate and in monolayer, an additional 150 µL of RPMI medium was added. After 24-48 hours, cell plates were washed twice with pre-warmed Seahorse medium, following the manufacturer’s instructions for each assay type (Seahorse, Agilent Technologies, St. Clara, CA, United States), and incubated at 37°C in the oven for 1 hour. In parallel, the calibration cartridges were loaded into the equipment Seahorse XFe24 Analyzer (Seahorse, Agilent Technologies, St. Clara, CA, United States) with the appropriate treatments and after 20 minutes cell plates were loaded for 1-2 hours. All experiments were performed in triplicate for each cell line and the results obtained were analyzed on the manufacturer’s own website (Seahorseanalytics.agilent).

### TMA, immunohistochemistry, and quantification

Tissue microarrays (TMA) containing 69 cores of nevus (n=14), melanoma (n=27) or metastasis (n=28) samples, were constructed using the MTA-1 tissue arrayer (Beecher Instruments, Sun Prairie) for immunohistochemistry analysis. Each core (diameter 0.6 mm) was punched from pre-selected tumor regions in paraffin-embedded tissues. We chose central areas from the tumor, avoiding foci of necrosis. Staining was conducted in 2-μm sections. Slides were deparaffinized by incubation at 60°C for 10 min and incubated with PT-Link (Dako, Agilent) for 20 min at 95°C in low pH to detect PGM1, PYGB and PYGL or in high pH to detect GYS. To block endogenous peroxidase, holders were incubated with peroxidase blocking reagent (Dako, Agilent) and then with (1:50) dilutions of antibody anti-PGM1, (1:25) anti-GYS, (1:100) anti-PYGB, and (1:150) anti-PYGL, overnight at 4°C. All previously described antibodies presented high specificity. After that, slides were incubated for 20 min with the appropriate anti-Ig horseradish peroxidase-conjugated polymer (EnVision, Dako, Agilent). Sections were then visualized with HRP Magenta (Dako, Agilent) as a chromogen for 5 min and counterstained with Harrys’ Hematoxylin (Sigma Aldrich, Merck). Photographs were taken with a microscope (Zeiss) with a 20x objective. Immunoreactivity was quantified blind with a Histoscore (H score) that considers both the intensity and percentage of cells stained for each intensity (low, medium, or high) following this algorithm (range 0–300): H score = (low%) × 1 + (medium%) × 2 + (high %) × 3. Quantification for each patient biopsy was calculated blindly by 2 investigators (MJFA and JMU). Clinicopathological characteristics of patients are summarized in Table 1.

### RNA-seq

STR authenticated, mycoplasma free (Lonza #LT07-318, #LT07-518 (control) melanoma cell lines were seeded in 6 well tissue cultured dishes in RPMI 1640 (Gibco) supplemented with 10% FBS (Biosera) and 100 U/ml Pen Strep (Gibco) and maintained in 10% CO2 humidified chamber. Cells were collected at around 80% confluence and snap-freeze on dry ice before batched processed using rNeasy Mini Kit (QIAGEN #74106) as per supplier instructions and eluted in 50 µl nuclease free water (Invitrogen #10977049). Samples with RIN values ≥9.5 (assessed using Agilent RNA 6000 Nano Kit (Agilent #5067-1511)) were carried forward for library prep using QuantSeq Forward kit (LEXOGEN #015.96) with 500 ng input material and ERCC ExFold RNA Spike-In Mixes (ThermoFisher #4456739). Sequencing was carried out on a HiSeq4000 (Illumina) by the Oxford Genomics Centre, Wellcome Trust Centre for Human Genetics.

### Bioinformatics

Raw fastq reads for the same samples sequenced across 2 lanes were stitched using UNIX, qCed using fastqc (v0.11.9), adaptor- and poly-A- trimmed using Cutadapt. Processed fastq were mapped against hg38 (GRCh38, 2015) + ERCC STAR index using rna-star (v2.5.1b) with quantMode enabled, allowing for soft-clipping and splicing (min 20 bp). Normalisation and differential gene expression analyses was performed as previously described (11, 102, 103) using edgeR glmQLFTest. Heatmaps were visualised using R package pheatmap (v1.0.12).

Gene expression from 53 melanoma cell lines (30) were obtained from GSE80829 and their classification as per the original publication.

Gene expression data from the CCLE and TCGA were accessed through cBioportal (104, 105). Moving average trendline of bar plot representing expression level of individual samples was generated using a simple window size of 20. Spearman correlation and the significance was calculated using cor.test function in R.

Gene expression from samples of melanoma and healthy tissue adjacent to the tumor were analyzed with TNMplot.com (https://tnmplot.com/analysis/) tool 59. Kaplan-Meier curves performed using the best cut-off point and data from 98 melanoma patients from the The Human Protein Atlas (https://www.proteinatlas.org/).

### Statistical analysis

Results are presented as fold induction, mean ± SEM from at least three biological replicates. Comparisons between two independent groups were performed with Student’s t test. Tests for significance between two sample groups were performed with Student’s t test and for multiple comparisons, ANOVA with Bonferroni’s post-test. All statistical analyses were performed with GraphPad Prism software. P-values < 0.05 were considered statistically significant.

For immunohistochemical data, the Kolmogorov-Smirnov test was used to determine whether calculated H- scores for each of the antigens were well-modelled by a normal distribution. None showed normal distribution. Analyses between groups were performed with Mann-Whitney U-test and correlations with Spearman test (S). All statistical analyses were performed with SPSS IBM software v.24. P-values < 0.05 were considered statistically significant.

## Ethics Statements

Human samples were obtained from Hospital Clínico San Carlos (HCSC). IRB-HCSC act number 21/498-E granting approval on June 25^th^, 2021.All patients gave written informed consent for the use of their biological samples for research purposes. Fundamental ethical principles and rights promoted by Spain (LOPD 15/1999) and the European Union EU (2000/C364/01) were followed. In addition, all patients’ data were processed according to the Declaration of Helsinki (last revision 2013) and Spanish National Biomedical Research Law (14/2007, of July 3).

## Acknowledgements

Funding was provided by Agencia Estatal de Investigación MCIN/AEI/10.13039/501100011033; PID2021- 127645OA-I00 (A.C-C) and PID2023-151128OB-I00; PID2019-104867RB-I00 (C.G.J.); and Comunidad de Madrid PRECICOLON-CM, P2022/BMD-7212 (C.G.J.) and Ayudas Atracción de Talento 2017-T1/BMD- 5334 and 2021-5A/BMD-20951 (A.C-C). This work was also supported by the Ludwig Institute for Cancer Research (CRG, PL), The National Institutes of Health R01 CA268597-01 (C.R.G., P.L.).

## Author Contributions

A.C-C and CG-J. conceived the project and designed and interpreted experiments. A.C-C., A.R-S, M.J-O., J.M. G-M, I. T-M and T. O-D performed experiments, C.R.G. and P.L. provided bioinformatic analysis, J. M- U and MJ. F-A provided melanoma samples and undertook IHC analysis. A.C-C and C.G-J. provided resources and supervision. A.C-C., C.R.G. and CG-J. wrote the manuscript.

---

**Fig. S1:**
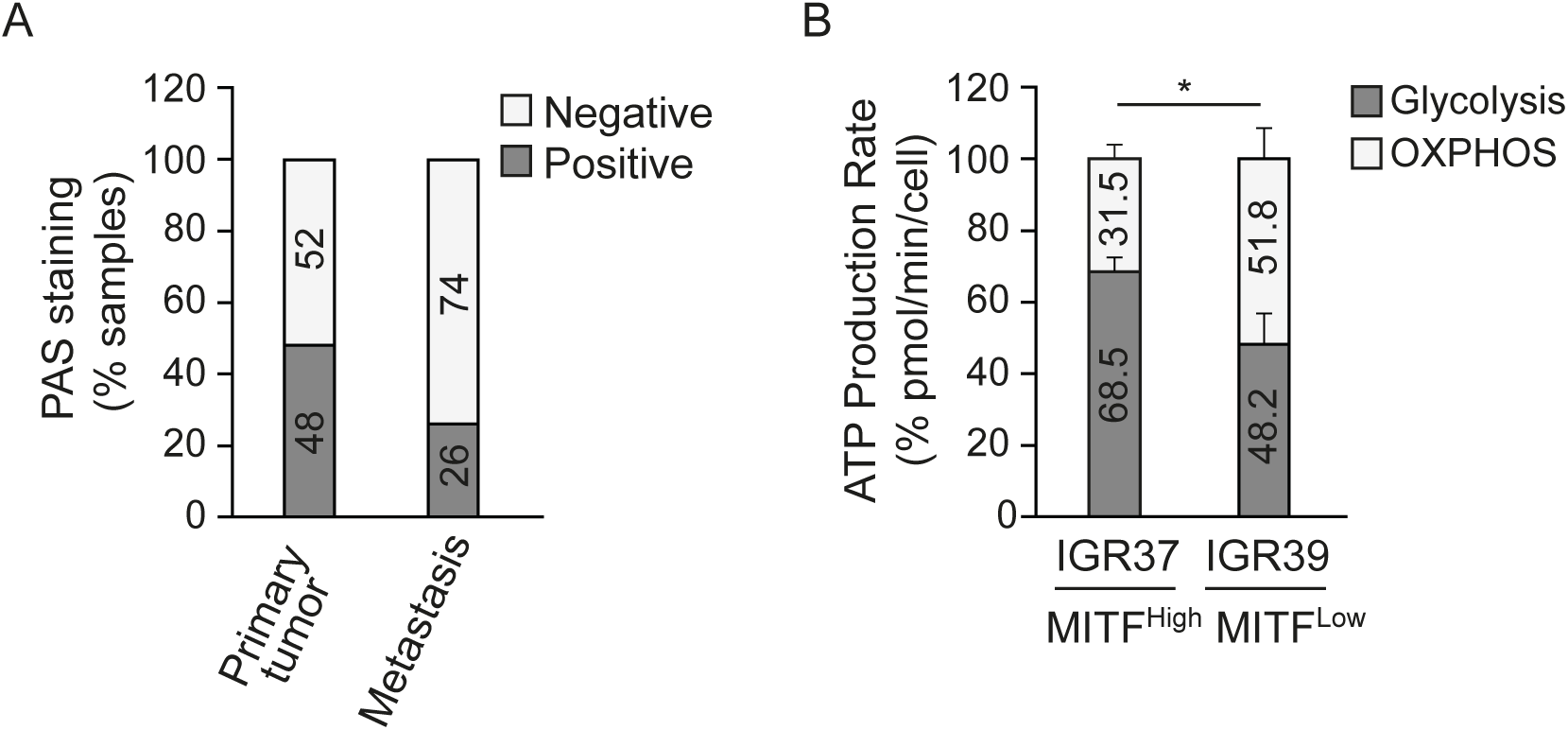
(A) Graphic representation of the percentage (%) of samples that store glycogen (PAS Positive) and that do not store glycogen (PAS Negative) in melanoma and metastasis biopsies. (B) Percentage of ATP production rate from glycolysis or mitochondrial oxidative phosphorylation (OXPHOS), for MITF^High^ (IGR37) and MITF^Low^ (IGR39) melanoma cells. n=4; error bars indicate SEM * P<0.05. Paired t test.

**Fig. S2:**
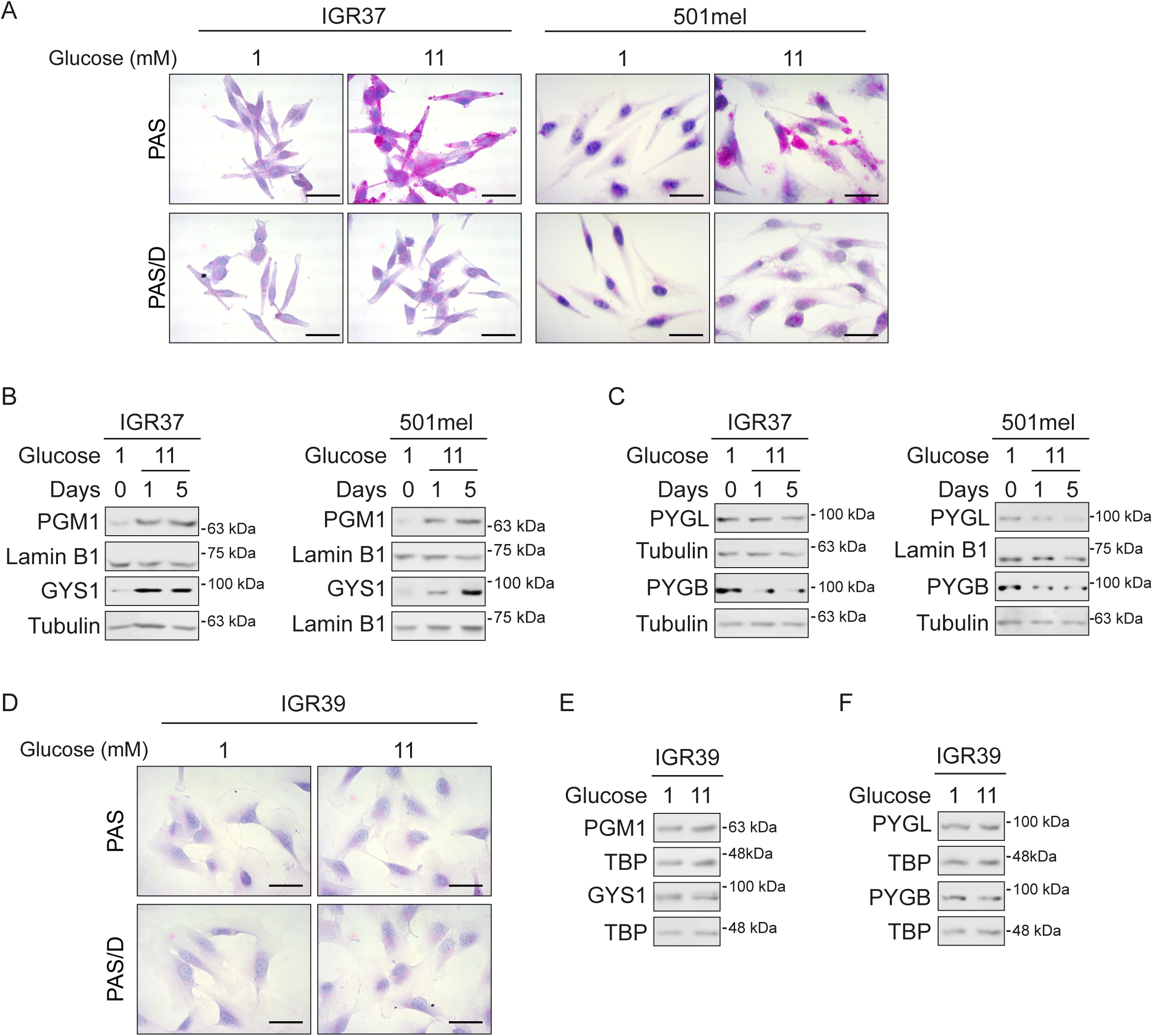
(A) Glycogen content of IGR37 and 501mel melanoma cells growing in 1mM glucose for 24h and treated with 11mM for another 24h. Digestion with diastase (PAS/D) is used to assess the specific glycogen staining. Scale bars: 25μm. (B-C) Western blot showing expression of indicated proteins in IGR37 and 501mel (melanoma cells growing as A and treated with 11mM for 1 or 5 days. Tubulin or lamin B are used as loading control. (D) Glycogen content of IGR39 melanoma cells growing as in A. (E-F) Western blot showing expression of indicated proteins in IGR39 treated as in D. TBP is used as loading control.

**Fig. S3:**
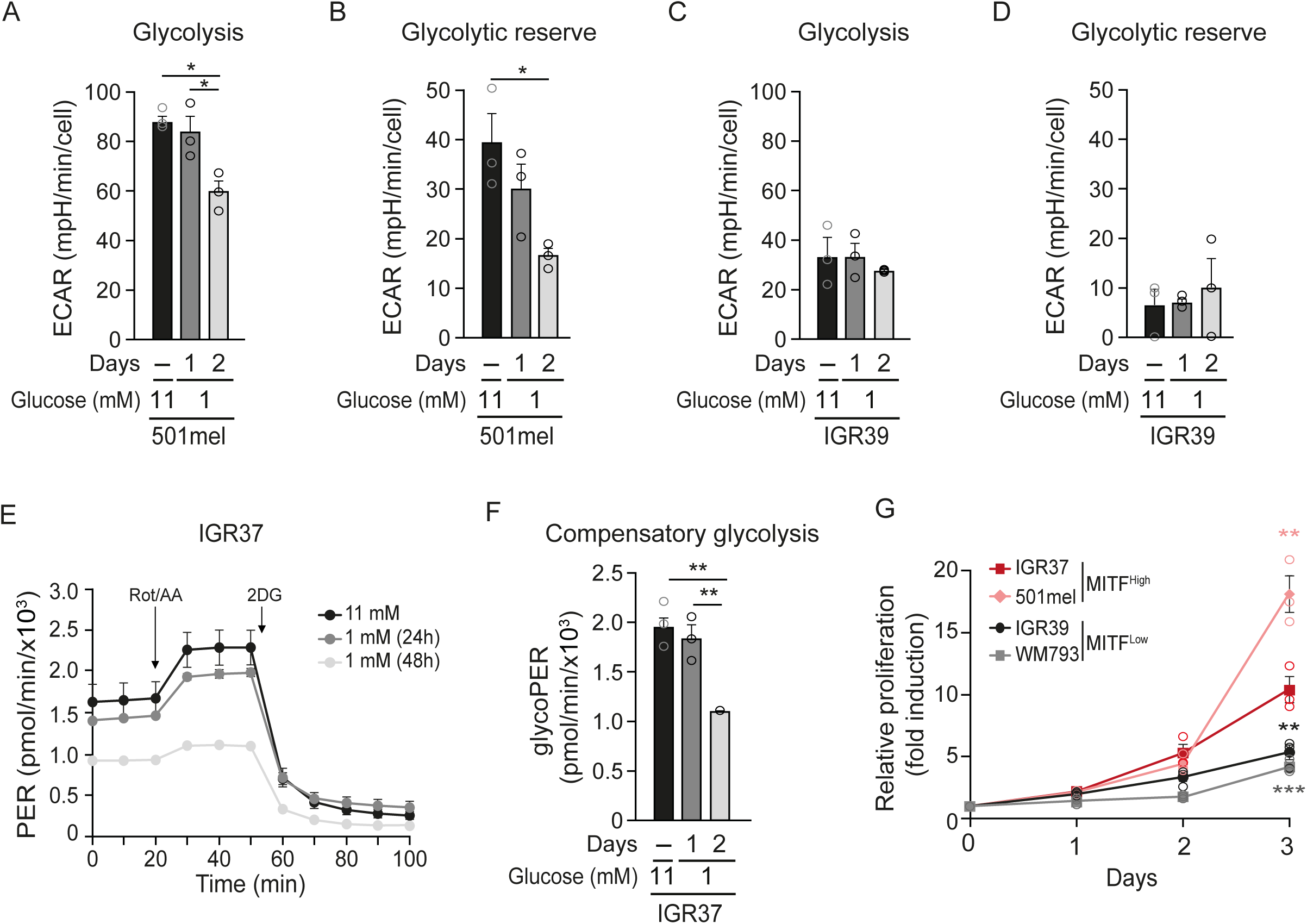
(A-C) Quantification of Basal glycolysis (A-C) and the glycolytic reserve (B-D) measurement derived from the Extracellular acidification rate (ECAR) of 501mel (A-B) and IGR39 (C-D) melanoma cells. n=3; error bars indicate SEM * P<0.05, One-Way ANOVA Statistical test. (E) Glycolytic Proton Efflux Rate profile (PER) from IGR37 melanoma cells depleted for glucose (1mM) for 1 and 2 days. The media of 3 experiments is shown; error bars indicate SEM. (F) Quantification of compensatory glycolysis measurement derived from D. n=3; error bars indicate SEM * P<0.05, One-Way ANOVA Statistical test. (G) Basal proliferation of MITF^High^ (IGR37, 501mel) and MITF^Low^ (IGR39, WM793) melanoma cells growing in 11mM glucose for 3 days. Statistical analysis by t-Student; n=3; error bars indicate SEM * P<0.05, **p < 0.01, ***p < 0.001. One- Way ANOVA Statistical test.

**Fig. S4:**
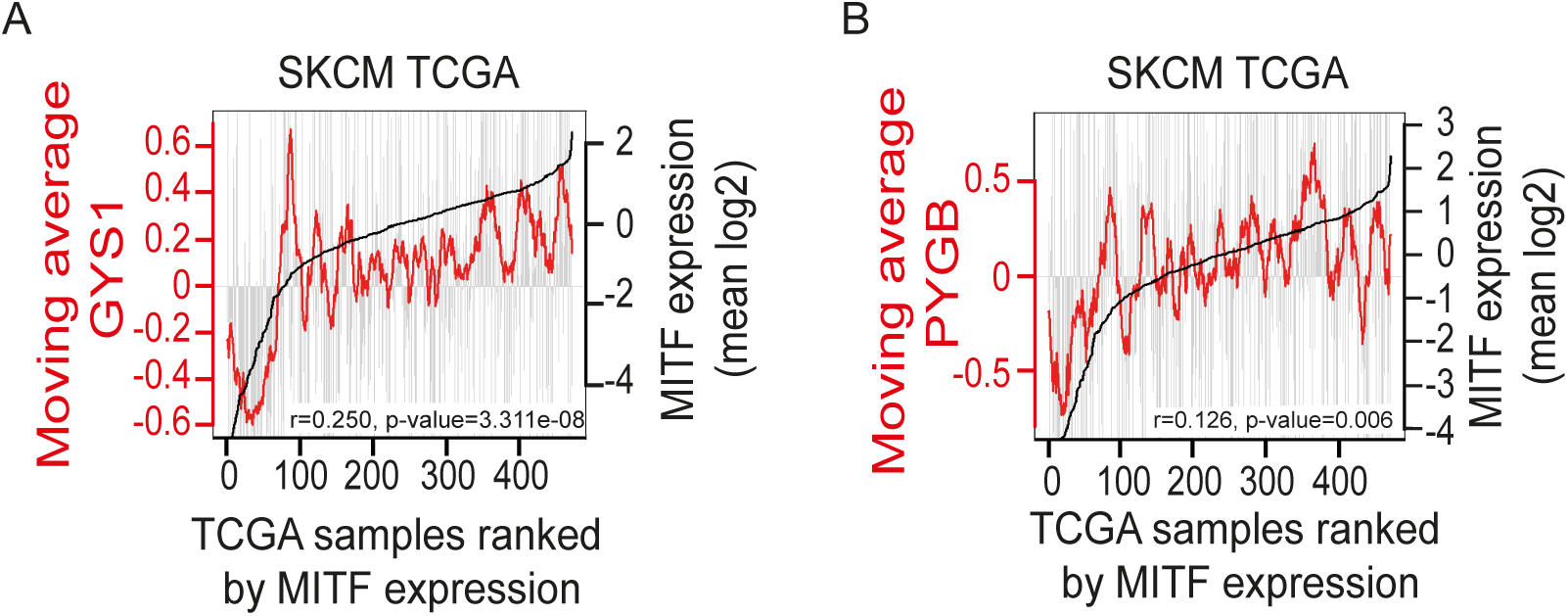
(A-B) Plots showing TCGA melanomas ranked by expression of MITF (Black lines). Expression of GYS1 (A) or PYGB (B) in each melanoma is indicated by grey bars, and the moving average of each per 20 melanoma window is indicated by the coloured lines. P value and correlation coefficient (rho) are indicated.

**Fig. S5:**
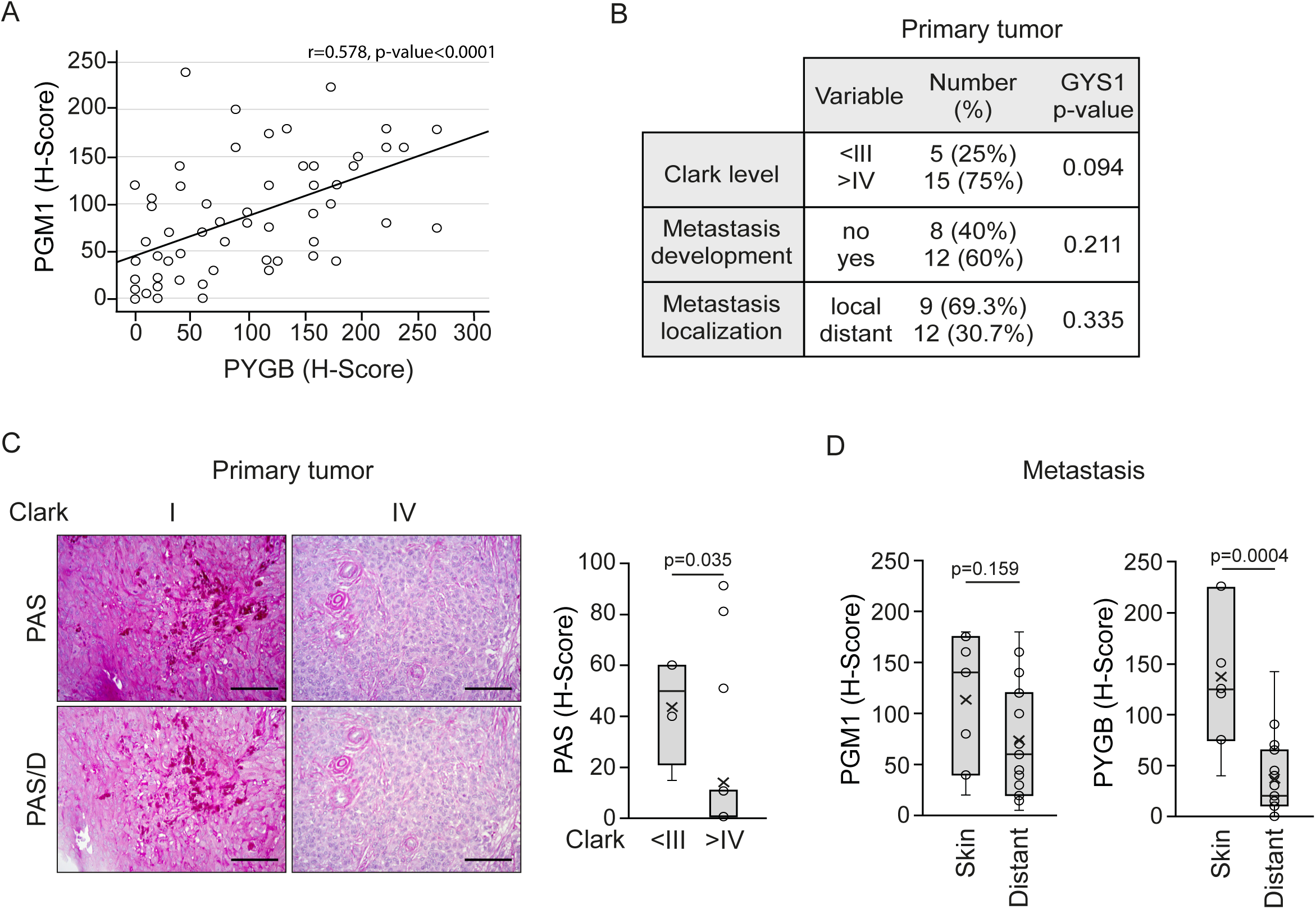
(A) Spearman rank correlation between PGM1 and PYGB protein levels on human melanoma samples. Correlation Rho (R) and p value are shown. ***p < 0.001. (B) Association analysis between GYS1 and clinical-pathological characteristics. Statistical analysis by Man Whitney U-test. The statistical significance cut off was set at p<0.01. (C) Representative images of glycogen content of human melanoma biopsies at I or IV Clark level stained with PAS or digested with diastase (PAS/D). Scale bars: 50 μm. Profile for Glycogen (PAS) content from melanoma human samples is shown. The profile was obtained from Histoscore variations between melanomas with high Clarck level (>IV) or low level (<III). Statistical analysis by Man Whitney U-test; *p<0.05, **p<0.01 and ***p<0.001. (C) Profile for PGM1 or PYGB content from melanoma metastasis samples ranked by the metastasis localization, skin or distant metastasis (ganglion, lung or hepatic) localization (D). Statistical analysis by Man Whitney U-test. *p < 0.05.

